# Permutational immune analysis reveals architectural similarities between inflammaging, Down syndrome and autoimmunity

**DOI:** 10.1101/2021.09.13.460115

**Authors:** Katharina Lambert, Keagan G. Moo, Azlann Arnett, Gautam Goel, Kaitlin J. Flynn, Cate Speake, Alice E. Wiedeman, Carla J. Greenbaum, S. Alice Long, Rebecca Partridge, Jane H. Buckner, Bernard Khor

## Abstract

People with Down syndrome show cellular and clinical features of dysregulated aging of the immune system, including naïve-memory shift in the T cell compartment and increased incidence of autoimmunity. However, a quantitative understanding of how various immune compartments change with age in Down syndrome remains lacking. Here we performed deep immunophenotyping of a cohort of individuals with Down syndrome across the lifespan, selecting for individuals not affected by autoimmunity. We simultaneously interrogated age- and sex-matched healthy neurotypical controls and people with type 1 diabetes, as a representative autoimmune disease. We built a new analytical software, IMPACD, that enabled us to rapidly identify many features of immune dysregulation in Down syndrome that are recapitulated in other autoimmune diseases. We found significant quantitative and qualitative dysregulation of naïve CD4^+^ and CD8^+^ T cells in Down syndrome and identified IL-6 as a candidate driver of some of these changes, thus extending the consideration of immunopathologic cytokines in Down syndrome beyond interferons. Notably, we successfully used immune cellular composition to generate three quantitative models of aging (i.e. immune clocks) trained on control subjects. All three immune clocks demonstrated significantly advanced immune aging in people with Down syndrome. Notably, one of these clocks, informed by Down syndrome-relevant biology, also showed advanced immune aging in people with type 1 diabetes. Together, our findings demonstrate a novel approach to studying immune aging in Down syndrome which may have implications in the context of other autoimmune diseases.

**One Sentence Summary:** Permutational analysis of immune landscape reveals advanced immune aging in people with Down syndrome and in people with type 1 diabetes.

## INTRODUCTION

Down syndrome (DS) is caused by trisomy 21 and is the most common congenital chromosomal abnormality, affecting about 1 in 800 births *(1)*. Immune dysregulation is a central feature of DS, manifested clinically by a greatly increased predisposition to many autoimmune diseases including autoimmune thyroid disease, type 1 diabetes (T1D), celiac disease and psoriasis *(2, 3)*. Autoimmunity is an important and growing burden on quality of life in people with DS due in part to the increasing lifespan of people with DS increasing the duration of this chronic comorbidity *(2)*. Understanding how triploidy of the 222 genes on chromosome 21 remodels the immune landscape is essential to enable rational selection and development of therapies to specifically mitigate DS-associated autoimmunity *(4)*. Furthermore, this understanding can advance our understanding of how related mechanisms promote autoimmunity in the broader (neuro)typical (i.e. people without DS) population.

Previous studies have identified several features of immune remodeling in DS associated with autoimmunity, including altered immune cellular subsets (e.g. increased memory:naïve T cells), elevated circulating levels of pro-inflammatory cytokines and transcriptional evidence of a hyper-response to type 1 interferons *(5-10)*. Interestingly, many of these features have also been described in the context of inflammaging, an inflammation-driven remodeling of the immune landscape during aging associated with increased risk of autoimmunity *(11-13)*. The role of accelerated immune aging has not been well-studied in the context of DS. It remains to be clarified (i) how DS impacts immune remodeling across the lifespan, (ii) what features of immune remodeling are DS-specific, (iii) the role of type 1 interferon and other cytokines in DS-immune remodeling, and (iv) how autoimmunity impacts immune remodeling in DS. Prior immunologic studies frequently focus on either pediatric or adult subjects with DS, hampering a comprehensive study of immune changes across the lifespan. Additionally, many DS cohorts are enriched for autoimmunity, which may influence immune age *(14)*. Therefore, we lack to date a quantitative understanding of how immune age is altered in DS, when during the lifespan immune age differences begin and whether these differences contribute to the development of comorbidities or vice versa.

In this study, we addressed these questions by evaluating the immune landscape in a cohort of individuals with DS, ages 2 to 55 years old, with a relatively low incidence of autoimmunity. We included age- and sex-matched healthy typical control participants, as well as typical individuals with T1D to identify changes related to one autoimmune disease across a wide range of ages. We built a new software tool, IMPACD (Iterative Machine-assisted Permutational Analysis of Cytometry Data), to improve analytic rigor of manual gating analyses and perform exhaustive permutational analysis of mass cytometry data with zero down-sampling. Using IMPACD, we found extensive dysregulation of naïve T cells in DS and identified IL-6 as a potential driver of several of these qualitative changes. We also found evidence of co-regulation of CD4^+^ and NKT cells in DS. Importantly, IMPACD empowered us to build immune clocks using immune subsets that correlate linearly with age in typical controls. These immune clocks quantitatively demonstrate advanced immune aging in DS for the first time to our knowledge. Furthermore, a DS-informed immune clock also demonstrated advanced immune aging in people with T1D. These findings advance our understanding of immunopathology in people with and without DS.

## RESULTS

To investigate features of immune dysregulation in DS that may underlie predisposition to autoimmunity, we used mass cytometry (CyTOF) to immunophenotype PBMCs from 28 individuals with DS (ages 2 to 55 years), 28 age- and sex-matched healthy typical controls (TCs) and 25 age- and sex-matched typical individuals with T1D (n=28) (Fig. 1A-B). Autoimmunity in our DS cohort was limited to Hashimoto’s disease, a very common cause of hypothyroidism, in 2 of 28 participants. As CMV infection can significantly alter the immune landscape, we measured anti-CMV antibodies as evidence of prior infection and found similar proportions of seropositive subjects in all cohorts (Fig S1A) *(15)*.

**Fig. 1.**
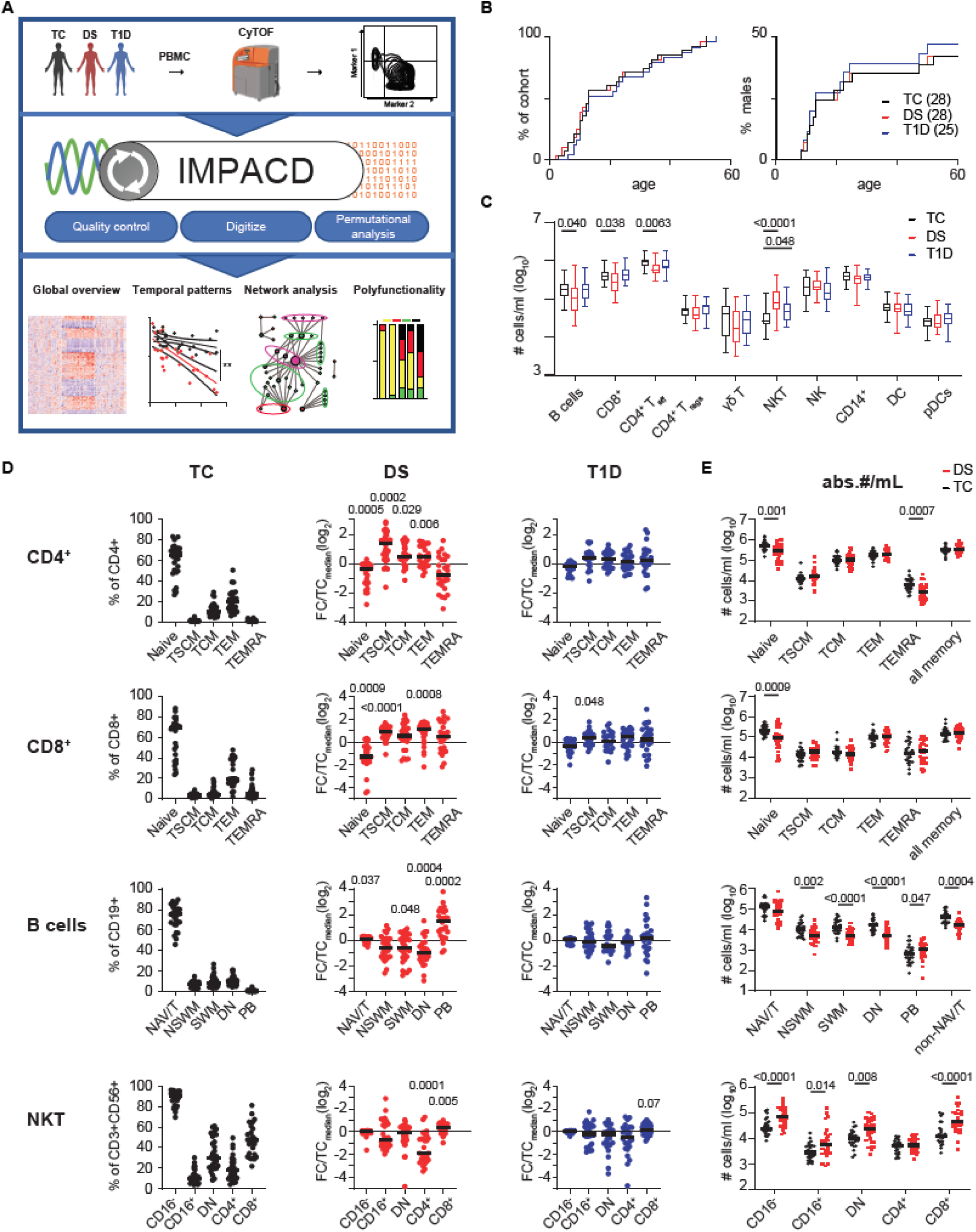
Broad changes in immune architecture in Down syndrome revealed by mass cytometry. (**A**) Schematic of study design. PBMCs from age and sex-matched participants with Down syndrome (DS, n=28), typical controls (TC, n=28) or typical participants with T1D (T1D, n=25) were immunophenotyped by mass cytometry. A new software tool, IMPACD, was developed to perform rigorous and exhaustive permutational analysis. IMPACD’s rich data output is readily analyzed using ‘omics-relevant approaches. (**B**) Age and sex distribution of cohort. (**C**) Absolute numbers of major immune cell types, by cohort. (**D**) Frequencies of CD4^+^ T, CD8^+^ T, B cell and NKT subsets in TC, DS and T1D participants. Data from participants with DS or T1D is scaled (FC, fold change) to the corresponding subset’s median value in TC participants (TC_median_). (**E**) Absolute numbers of CD4^+^ T, CD8^+^ T, B cell and NKT subsets in TC and DS participants. (**B-E**) n = 28 (**B, D**) or n = 26 (**C, E**) TC, n = 28 DS, n = 25 T1D, across 5 batches. Mann-Whitney test, p-values shown.

### Broadly altered adaptive immune landscape in participants with Down syndrome

We first quantitated proportions of major innate and adaptive cell types by manual gating and absolute numbers using contemporaneous complete blood count (CBC) data. We found decreased absolute numbers of B cells, CD4^+^ T_eff_ and CD8^+^ T cells, consistent with previous studies showing varying degrees of B- and T-lymphopenia in DS (Fig. 1C and Fig. S1B) *(7, 10, 16)*. Numbers of T_regs_ were comparable between cohorts (Fig. 1C). CD3^+^CD56^+^ NKT cells were increased in DS and, to a lesser extent, T1D (T1D, Fig. 1C). Analyses of cellular frequencies among total PBMCs showed similar trends (Fig. S1C).

As T and B cells were the most quantitatively altered subsets in DS, we next assessed qualitative remodeling in these cells, first enumerating naïve and memory subsets as previously defined *(17)*. Given the wide range of frequencies between these subsets, we show the raw percentage in TC and normalize the percentage in people with DS or T1D to the corresponding median value in TC to effectively display all data simultaneously (Fig. 1D). Consistent with previous studies, we found decreased frequencies of naïve CD4^+^ T_eff_ and CD8^+^ T cells in DS (Fig. 1D and Fig. S1B) *(7, 10)*. The frequency of most memory T cell subsets in DS was increased, including stem cell memory (SCM) and effector memory (EM) T cells (Fig. 1D and Fig. S1B). Central memory (CM) cells were increased only in CD4^+^ T_effs_; TEMRAs were increased in neither CD4^+^ T_eff_ nor CD8^+^ T cells (Fig. 1D and Fig. S1B). We did not observe any differences in frequencies of naïve or memory cells amongst T_regs_ (Fig. S1D). We calculated the absolute number of cells in each subset and found that naïve CD4^+^ and CD8^+^ T cells were significantly decreased without any increase in any memory T cell subset; in fact, CD4^+^ TEMRAs were decreased in DS (Fig. 1E). These results show that the increased memory:naïve T cell ratio is driven by decreased absolute numbers of naïve T cells in DS, consistent with prior studies suggesting decreased thymic output of naïve CD4^+^ T cells in people with DS *(18)*.

In contrast, the DS-B cell compartment showed increased proportion of naïve B cells and decreased proportion of most memory subsets with the exception of plasmablasts (PB), which were increased (Fig. 1D). Enumerating absolute numbers revealed that TC and DS individuals have similar numbers of naïve B cells, but memory B cell subsets, except plasmablasts, are decreased in people with DS (Fig. 1E). These results demonstrate that decreased numbers of memory B cells drive the memory-naïve B cell shift in DS. Previous studies showing intact proliferation and somatic hypermutation in memory B cells in DS suggest that these differences reflect impaired T cell help, although B-cell-intrinsic dysregulation cannot be excluded *(19, 20)*.

We further examined NKT cells using two well-established classification schemes of CD16 or CD4/CD8 expression *(21, 22)*. We found a decreased proportion of CD4^+^ NKT cells that was driven by increased absolute numbers of CD8^+^ and CD4^-^CD8^-^ (DN) NKT cells (Fig. 1D-E). Previous studies demonstrating increased cytotoxic capacity of CD8^+^ and DN NKT cells suggests an immunomodulatory role *(23, 24)*.

Dysregulation of T, B and NKT compartments in T1D was less profound than in DS and included increased frequency of CD8^+^ SCM T cells and a trend (p=0.07) towards increased frequency of CD8^+^ NKT cells (Fig. 1D). Together, these results extend prior studies showing CD8^+^ SCM cells are important to T1D pathobiology and provide a unified view of major lymphocyte subset alterations in people with DS or T1D *(25, 26)*.

### IMPACD: High-rigor manual gating with exhaustive permutational analysis enhances deep immune subset profiling

We hypothesized that extending our analysis to encompass all markers would reveal important differences related to DS. However, the large number of markers (>40 in CyTOF) challenges the consistency of manual gating analysis of high-dimensional cytometry data *(27)*. One challenge is threshold variability, where graphical thresholding of one parameter in multiple plots leads to inadvertent if slight variation (Fig. S2A). Another challenge is “versioning”, which arises from the need to adjust and reapply gates to multiple subsets. To overcome these challenges, we developed a novel software tool, IMPACD, that reads and displays gate values from standard WSP files. This allowed us to easily identify and harmonize discrepant threshold values within and between batches (Fig. S2A). Infrequent use of batch-specific threshold values was guided by the internal control. Thus, IMPACD maximizes the consistency and rigor of manual gating analysis. Next, we used IMPACD to read standard FCS files, using the harmonized thresholds to binarize each cell as positive or negative for each marker (Fig. S2A). IMPACD and Flowjo counted very similar absolute numbers of cells as positive for each of 29 markers in all 81 subjects over a wide dynamic range, validating IMPACD’s performance (Fig. S2B).

We then used IMPACD to define differences in the immune landscape of DS, T1D and TC. Building on “root nodes” of biologically important T, B and NKT subsets described above, we exhaustively interrogated all permutations of up to three additional markers (Fig. 2A). We found diminishing yield of examining more markers, which mostly generated rare subsets (Fig. S2C). This approach allows us to map novel differences onto a scaffold of known biology (e.g. cell type/developmental stage). Each subset is described as a “path” of serially queried markers comprising the “root node”, 0-2 “modifier nodes” and the “terminal node” (Fig. 2A). Key features of our analysis include (i) zero down-sampling, (ii) rigorous multiple testing correction, (iii) exclusion of small (<10 cells in both cohorts) subsets and (iv) exclusion of uninformative “modifier nodes”, which significantly curates the output (Fig. 2B).

**Fig. 2.**
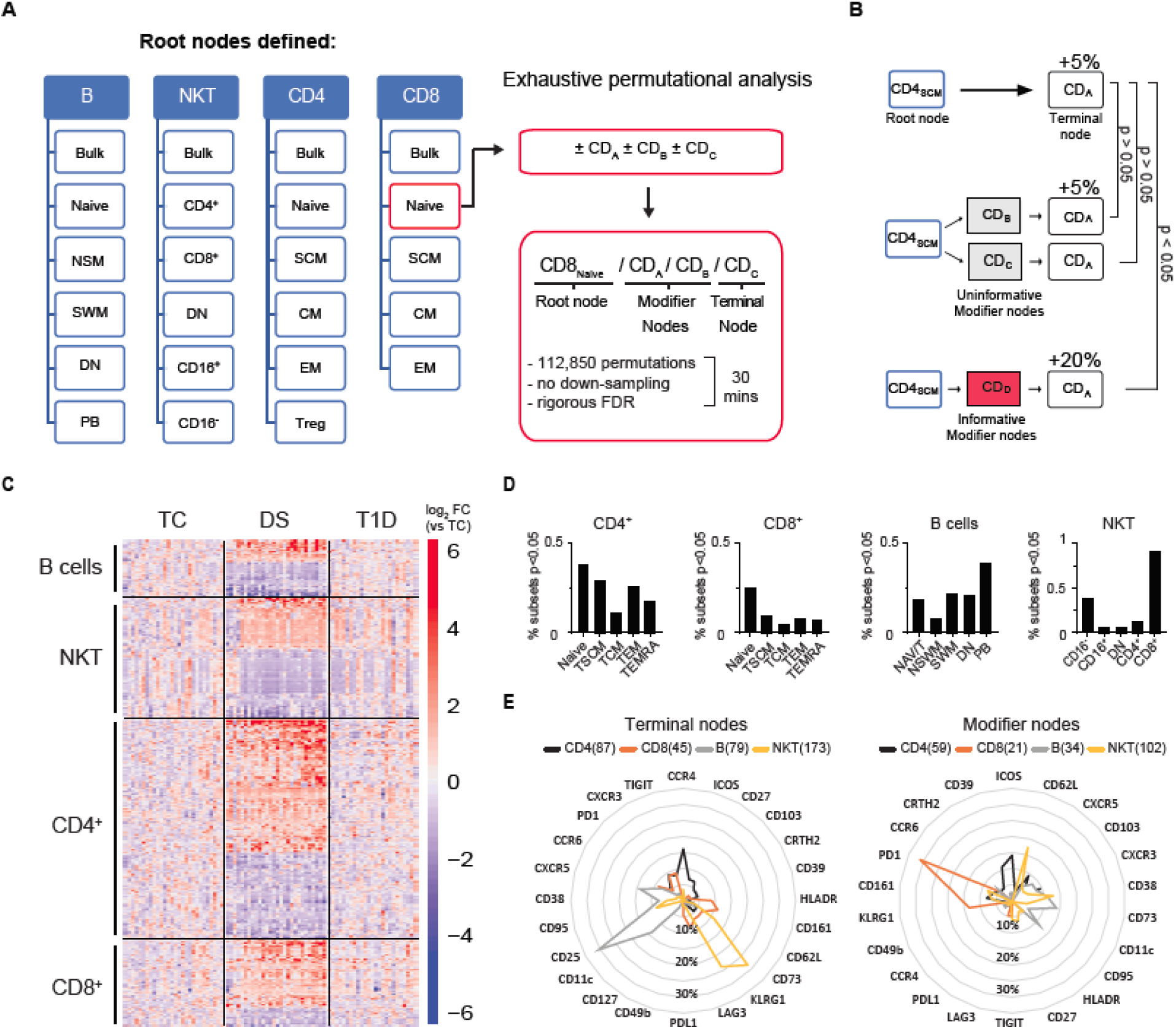
IMPACD performs deep immune subset profiling via permutational analysis. (**A**) IMPACD uses thresholds set by digital gating to identify defined subsets of interest and perform permutational analysis on each pre-defined subset using all remaining markers. Subsets are described by a path including root, modifier and terminal nodes as shown. (**B**) Outline of modifier analysis, which curates differentially represented subsets for maximal informativeness. In this hypothetical example, expression of CD_A_ is altered in CD4_SCM_ cells. Many subsets of CD4_SCM_ cells (e.g. CD4_SCM_CD_B_^+^, CD4_SCM_CD_C_^+^) would be expected to reflect this same difference and are curated out (gray boxes). IMPACD’s modifier analysis includes subsets that show statistically significant (Wilcoxon ranksum), additional differences (e.g. CD4_SCM_CD ^+^, red _D_box). *, p<0.05; ns, not significant. (**C**) Heatmap of B, T and NKT cell subsets identified by IMPACD as differentially abundant between control (TC) and DS and/or T1D. (**D**) Percentage of total identified subsets, in each cell type and memory subset, that were differentially abundant in DS versus TC (Wilcoxon ranksum test, nominal p<0.05). (**E**) Analysis of subsets differentially abundant in DS versus TC, comparing frequency of each modifier and terminal node in each cell type, as a proportion of all modifier/terminal nodes in that cell type.

### Qualitative immune landscape remodeling in Down syndrome

IMPACD analysis of B, T and NKT cells identified 651 subsets (i.e. combinations of expressed markers) that were differentially abundant in DS compared to TC (Fig. 2C). To reveal organizing principles, we categorized these 651 subsets according to cell type and differentiation stage. We found that dysregulated subsets in DS were non-uniformly distributed, with relative predominance in naïve CD4^+^, naïve CD8^+^, memory B cells, and CD8^+^ NKT cells (Fig. 2D). From this global viewpoint, qualitative changes tended to be concentrated in subsets that were also quantitatively altered in DS.

Next, we looked for common themes underlying these dysregulated subsets. First, we asked if similar markers were being dysregulated across different cell types by enumerating terminal nodes in each cell type. This showed that dysregulated terminal nodes exhibited highly cell-type-specific patterns with some limited overlap (Fig. 2E). Next, we assessed for commonalities between the subsets that exhibited dysregulated marker expression by enumerating modifier nodes in each cell type. We found that modifier nodes also exhibit cell-type-specific patterns with some limited overlap (Fig. 2E). The observation that each cell type exhibits dominant terminal and modifier nodes supports the notion that immune dysregulation in DS engages common mechanistic programs shared between subsets of each major cell type. The observation that different cell types exhibit different dominant terminal and modifier nodes could reflect regulation either by different effector mechanisms in each cell type or by common effectors modulated by cell-type-specific epigenetic landscapes.

### Qualitative remodeling of B and T cells in Down syndrome shows autoimmunity-related features

Dysregulated B and T cell homeostasis is a key feature of many autoimmune responses. To better understand similarities between B cell remodeling in DS and autoimmunity, we used IMPACD to build a temporal map of how B cells are dysregulated in DS at sequential maturation stages including naïve/transitional (NAV/T), non-switched memory (NSM), switched memory (SWM), double negative (DN) and plasmablasts (PB) (Fig. S1B and Fig. 3A). Qualitative remodeling of the B cell compartment in DS shared many features with other autoimmune diseases, including (i) increased expression of CD11c in non-PB B cell subsets, as seen in rheumatoid arthritis (RA), systemic lupus erythematosus (SLE) and multiple sclerosis (MS) *(28-30)*, (ii) decreased expression of CXCR5 and CD25 in non-PB B cell subsets, as seen in SLE and interestingly also in severe COVID-19 infection *(31, 32)*, (iii) increased expression of PD-1 in non-PB B cell subsets, as in RA *(33)*, (iv) increased CXCR3 and decreased CCR6 expression in SWM B cells, as in RA *(34)* and SLE *(35)*, (v) increased CD11c^+^CXCR5^-^ DN2 and CD11c^+^CXCR5^-^ activated naïve (aNAV) B cells, as in SLE *(31)* and (vi) increased expression of CD95 in DN B cells, as in SLE *(36)* (Fig. 3A-B). In contrast, the B cell compartment of people with T1D showed less profound remodeling (Fig. 3A). Together, these results highlight how the B cell compartment is broadly remodeled in DS to exhibit features consistent with several autoimmune diseases and impaired activation/function.

**Fig. 3.**
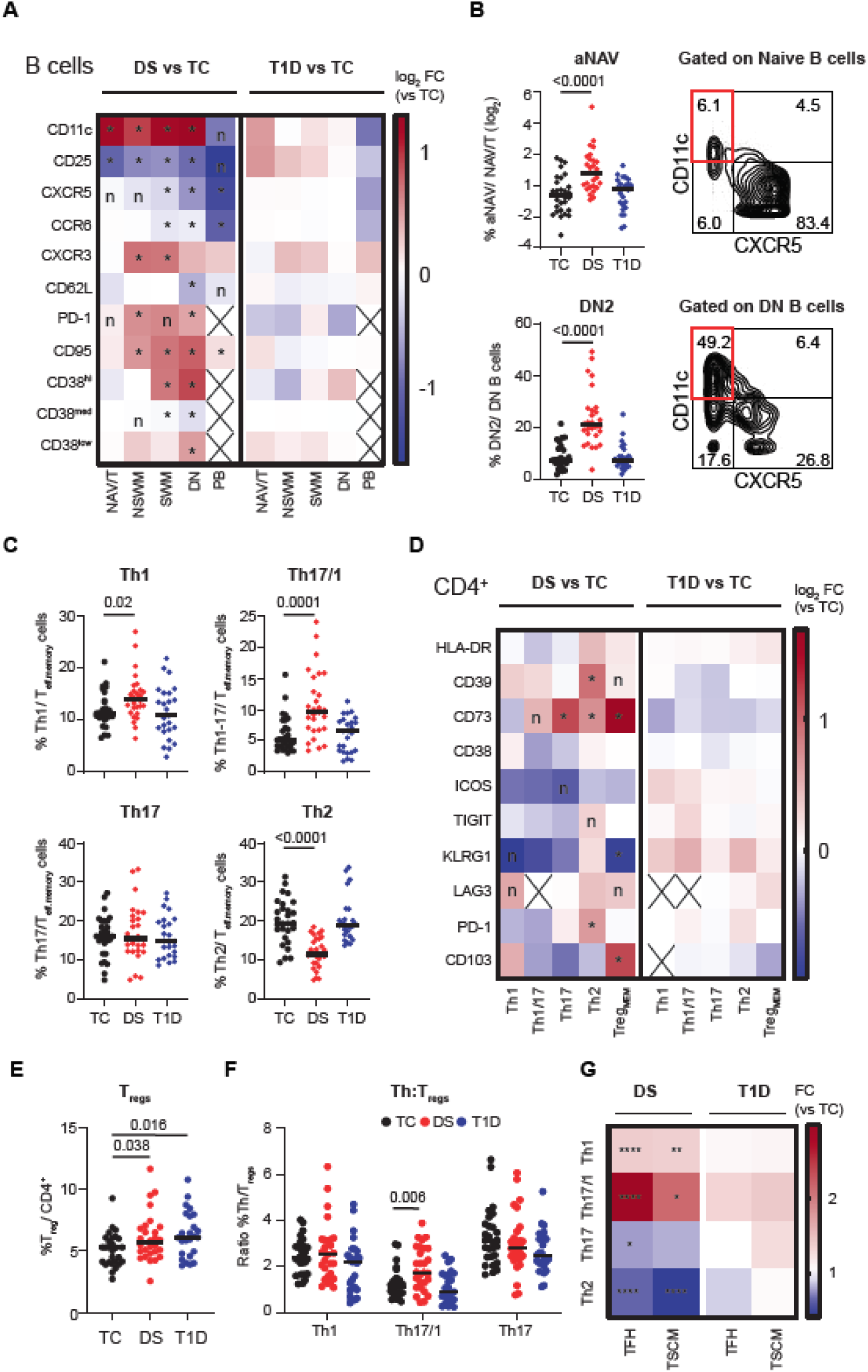
Remodeling of B and T cell compartments in people with DS shows autoimmunity-related features. (**A**) IMPACD compares abundance of each B cell memory subset, in either DS or T1D versus controls (TC), expressing each of several selected markers. Heatmap shows the ratio of median percentage of marker-expressing cells. NAV/T, naïve/transitional; NSWM, non-switched memory; SWM, switched memory; DN, double negative; PB, plasmablast. (**B**) Left, frequency of autoimmunity related activated naïve (aNAV) and DN2 B cells (amongst naïve and DN B cells respectively) in each cohort. Right, representative plots from a subject with DS showing gating of the respective subsets (red). (**C**) Frequencies of Th cell subsets among total non-T_FH_ (CXCR5^-^) CD45RO^+^ T_effs_ (T_eff.memory_) in each cohort. (**D**) Abundance of each Th subset, in either DS or T1D versus TC, expressing each of several selected markers. Heatmap shows the ratio of median percentage of marker-expressing cells. Treg_mem_, memory T_regs_. (**E**) Frequency of T_regs_ (CD127^low^CD25^hi^) amongst total CD4^+^ T_effs_ in each cohort. (**F**) Ratio of Th1, Th17/1 and Th17 cells to T_regs_ in people with DS or T1D or controls. (**G**) Frequency of Th subsets within total T_FH_ (CXCR5^+^CD45RO^+^ CD4^+^ T_eff_) or T_SCM_ cells from people with DS or T1D, normalized to median value in TCs. Heatmap shows median value. (**A**-**G**) n = 28 TC, n = 28 DS, n = 25 T1D, across 5 batches. (**A** and **D**) Wilcoxon ranksum test with Benjamini-Hochberg correction, *, FDR<0.05; n, nominal p<0.05; cross, median #cells<10. (**B, C, E**-**G**) Mann-Whitney test, p-values shown.

Similarly, we found that remodeling of T cells in DS also shared many features with other autoimmune diseases. Increased pro-inflammatory Th1, Th17 and Th17/1 subsets have been observed in MS *(37)*, Crohn’s disease *(38, 39)* and psoriasis *(40)*. Anti-inflammatory T_regs_ have been observed to be either increased or decreased in autoimmune diseases such as RA and SLE, suggesting complex regulation *(41-43)*. Based on surface expression of CXCR3, CCR6 and CCR4 (in CXCR5^-^ non-T_FH_ cells), we found increased Th1 and decreased Th2 frequency in DS (Fig. 3C). Although frequencies of Th17 cells were similar, the increase in Th17/1 cells, which are IFNγ^+^IL-17^+^ cells that arise from Th17 precursors, points to quantitative and qualitative remodeling of the Th17 compartment in DS (Fig. 3C) *(44)*. These results are consistent with previous reports of pro-inflammatory Th bias in DS *(6, 7)*. Next, we found that qualitative remodeling of surface markers was focused in Th2 and Th17 cells (Fig. 3D). Intracellular staining studies supported these findings by showing disproportionately increased expression of activation-associated cytokines (TNFα and IL-2) in IL-17^+^ Th17 cells (Fig. S3A). We found similarly increased frequencies of T_regs_ in both DS and T1D (Fig. 3E). Memory T_regs_ were also qualitatively remodeled specifically in DS, including increased expression of CD39 and CD73, resembling changes seen in the joints of patients with RA (Fig. 3D) *(45)*. The ratio of pro-inflammatory Th17/1 to anti-inflammatory T_regs_ was specifically increased in DS, highlighting Th17/1 cells as putative drivers of pathologic inflammation in DS (Fig. 3F). Finally, we examined two T cell subpopulations of particular interest in autoimmunity. T_FH_ cells help drive the B cell response and T_SCM_ cells can regenerate effector T cells, which may empower long-lived autoimmune responses *(46, 47)*. T_FH_ frequency was not significantly altered, but the frequency of both CD4^+^ and CD8^+^ T_SCM_ cells was increased in DS, as has been observed in RA *(48)* and aplastic anemia *(49)* (Fig. 1D and Fig. S3B). Both T_SCM_ and T_FH_ compartments are qualitatively remodeled in DS, showing similar skewing towards Th1- and Th17-like cells and away from Th2-like cells (Fig. 3G) *(50)*. Taken together, these results provide a detailed map of how Th subsets are dysregulated in a pro-inflammatory autoimmunity-relevant manner in DS.

### Naïve CD4^+^ T cells in Down syndrome exhibit a poised state driven in part by IL-6

Focusing on T cells, we found dysregulation in all stages of maturation, particularly in naïve CD4^+^ T_eff_ and naive CD8^+^ T, again showing qualitative remodeling focused on quantitatively altered subsets (Fig. 2D and 4A). Interestingly, naïve T cell changes involved many markers associated with T cell activation, including CD62L and CD38 (downregulated with T cell maturation) as well as CXCR3, TIGIT, KLRG1 and HLA-DR (upregulated with T cell activation) *(7, 51)*. Naïve CD4^+^ T_eff_ and CD8^+^ T cells from participants with DS showed overlapping but non-identical differences compared to TCs. Dysregulation of CD73, CD38 and ICOS was naïve CD4^+^ T_eff_-specific, upregulation of KLRG1 and HLA-DR was naïve CD8^+^ T cell-specific and decreased CD62L and increased CXCR3, CD39 and TIGIT was common to both naïve CD4^+^ T_eff_ and CD8^+^ T cells from people with DS (Fig. 4A). Changes in NKT cells broadly resembled the corresponding (CD4^+^ or CD8^+^) T cell subset (Fig. S3C). Remodeling in the setting of T1D was less profound (Fig. 4A). These findings suggest that naïve CD4^+^ and CD8^+^ T cells in people with DS exist in a hyperactivated state, poised for activation. This is further supported by our CyTOF findings of increased expression of intracellular activation-related markers including Ki-67, IL-2 and TNFα in naïve CD4^+^ T_eff_ and CD8^+^ T cells from people with DS, as well as previous studies showing enhanced effector differentiation potential of and cytokine expression by CXCR3^+^ naïve cells (Fig. S3D) *(7, 51, 52)*.

**Fig. 4.**
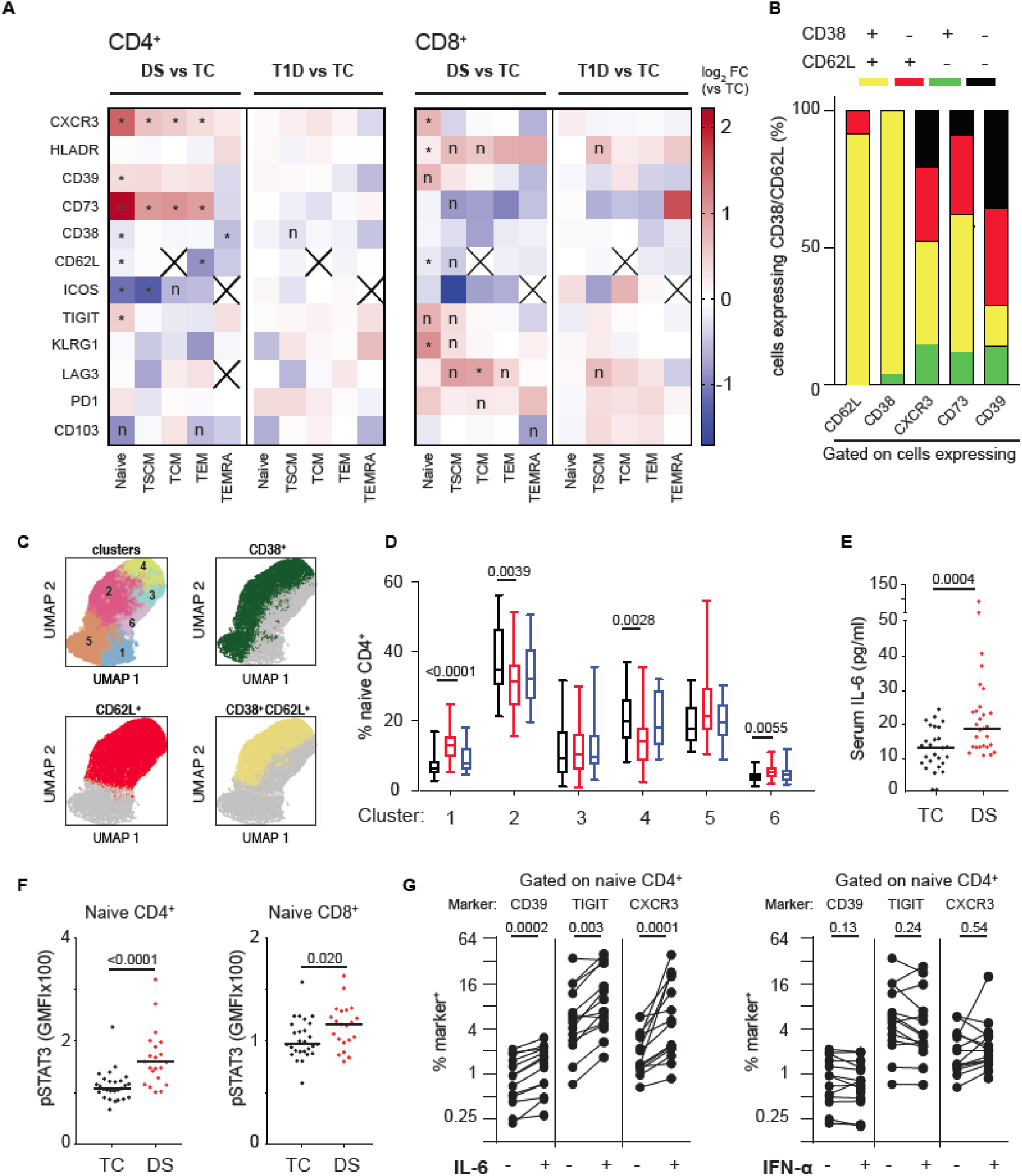
Remodeling of naïve T cells in people with DS suggests a state of poised activation driven in part by IL-6. (**A**) Abundance of each T cell memory subset, in either DS or T1D versus TC, expressing each of several selected markers. Heatmap shows the ratio of median percentage of marker-expressing cells. *, FDR<0.05; n, nominal p<0.05; cross, median #cells<10 or marker was part of definition (TCM). (**B**) Polyexpression analysis of selected markers dysregulated in naïve CD4^+^ T cells from people with DS shows significant co-expression of CD38 and CD62L (yellow). (**C**-**D**) FlowSOM analysis of naïve CD4^+^ T cells in TC, DS and T1D identifies 6 major clusters (upper left). Cells expressing CD38 (green), CD62L (red) or both (yellow), as determined by manual gating thresholds, are shown in separate overlays. (**D**) Percent of naïve CD4^+^ T cells in each FlowSOM cluster, by cohort. (**E**) Serum IL-6 is increased in people with DS. (**F**) Baseline pSTAT3 is increased in naïve CD4^+^ and CD8^+^ T cells from people with DS. (**G**) Expression of CD39, TIGIT and CXCR3 in naïve CD4^+^ T cells from controls is specifically increased by treatment with IL-6 and not IFNα. (**A**-**F**) n = 28 TC, n = 28 DS, n = 25 T1D, across 5 batches. (**A**) Wilcoxon ranksum test with Benjamini-Hochberg correction, *, FDR<0.05; n, nominal p<0.05; cross, median #cells<10 or part of subset definition (CM = CD62L^+^). (**D**) Quasibinomial logistic models with age as a covariate and Holm correction, p-values shown. (**E**-**F**) Mann-Whitney test, p-values shown. (**G**) n = 14 in 3 independent experiments, Wilcoxon ranksum test, p-values shown.

To better understand whether any of these markers were coordinately dysregulated, we performed a polyexpression (analogous to polyfunctionality) analysis of these markers in naïve CD4^+^ T cells. We found a striking decrease in CD62L^+^CD38^+^ double-positive cells in DS, highlighting that CD62L^+^ and CD38^+^ are coordinately decreased in naïve CD4^+^ T cells (Fig. 4B). This was supported by independent FlowSOM analysis, which grouped naïve CD4^+^ T cells into 6 major clusters with CD62L^+^CD38^+^ cells predominating in clusters 2 and 4 (Fig. 4C). Both of these clusters were decreased in DS (Fig. 4D). FlowSOM analysis also showed some increase in a CD62L^+^ and a CD38^-^CD62L^-^ cluster, although no other definitive markers were identified (Fig. 4C-D). Demonstrating the power of IMPACD’s downsampling-free and controlled-granularity approach, polyexpression analysis of CXCR3^+^, CD39^+^ and CD73^+^ cells revealed that the increased representation of these cells was associated with CD62L and/or CD38 co-expression (Fig. 4B). These results point to extensive remodeling of a CD62L^+^CD38^+^ compartment in DS, with a dramatic quantitative decrease accompanied by qualitative remodeling exemplified by upregulation of CXCR3, CD39 or CD73 expression.

To explore the role of cytokine signaling on the naïve T cell dysregulation in DS, we examined the response of TC T cells to IFNα and IL-6 in the absence of TCR stimulation. We focused on these cytokines because: (i) interferon signaling is enhanced in DS due to overexpression of chromosome 21-encoded interferon receptor subunits and supported by our findings of elevated phospho-STAT1 in DS T cells (Fig. S3E), and (ii) IL-6 is associated with autoimmunity, increased in serum from participants with DS and its functional relevance is supported by increased phospho-STAT3 in CD4^+^ T cells from people with DS (Fig. 4E-F) *(8, 9, 53, 54)*. Cytokine treatment did not deplete naïve T cells, supporting feasibility of our approach (Fig. S3F). Unexpectedly, we found that treating TC T cells with IL-6, but not IFNα, recapitulated many differences found in naïve CD4^+^ T cells from people with DS, including increased expression of CD39, TIGIT and CXCR3 (Fig. 4G).

### Identifying a novel coregulated CD4-NKT module in Down syndrome

To illuminate organizing principles underlying immune dysregulation in DS, we analyzed the 651 subsets differentially abundant in DS for modules that were coordinately regulated in DS differently from TC. We identified 145 pairs that were co-regulated in DS and showed either non-significant coregulation or opposite correlation in TC. We categorized pairs according to the cell types involved and found overrepresentation in the CD4^+^-NKT group (Fig. 5A). To validate these findings, we performed a simulation experiment where we randomly re-assorted subjects to DS or TC cohorts (10,000 iterations) before assessing differentially correlated subsets, generating an FDR-adjusted p-value and showing statistical significance in 7 of 10 groups of interactions (Fig. 5A).

**Fig. 5.**
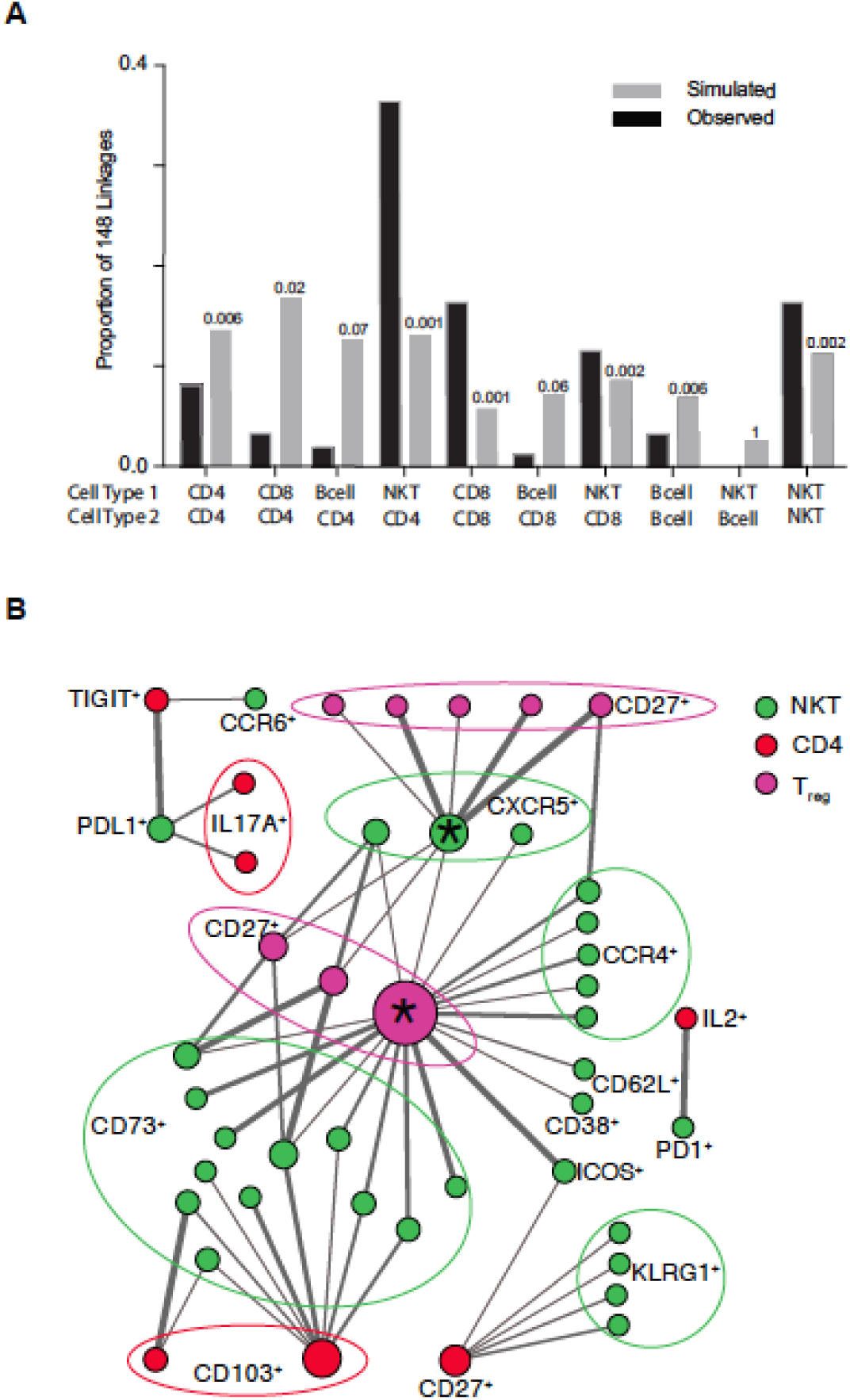
Identifying differentially co-regulated subsets in DS. (**A**) Diffcoex analysis of IMPACD data identifies subsets differentially co-regulated in DS versus TC, here organized by interacting cell types. For comparison, results from random choice simulations (x10,000) are shown in gray. Numbers above each column show the fraction of simulations where the number of differential correlations in each group reached or exceeded the same number as were actually observed. (**B**) Network map of CD4^+^ T-NKT interactions in DS reveals distinct organizational logic, including hubs. Two major inter-connected hubs are highlighted (*). Nodes are colored according to root node; size is proportional to number of connections. Terminal nodes are indicated, using boundary ellipses as appropriate. Edge thickness reflects significance of differential correlation in DS vs TC. (**A**-**B**) n = 28 TC, n = 28 DS, n = 25 T1D, across 5 batches.

We focused on the CD4^+^-NKT group, which had the highest significance and the largest number of co-regulated subsets (Fig. 5A and Table S1). We generated a network graph of the CD4^+^-NKT linkages and found distinct organizational logic. We further found that TIGIT^+^ naïve CD4^+^ T_eff_ were co-regulated with two NKT subsets specifically in DS (Fig. 5B and Table S1). We also uncovered larger patterns, including two inter-connected hubs, CD45RO^+^CD103^+^HLADR^-^CD27^+^ T_regs_ and CD95^+^CCR6^-^CXCR5^+^CD16^-^ NKTs (marked *) that were coordinately regulated with several (CD8^+^CD73^+^ and CD16^-^CCR4^+^) NKT and (CD27^+^) T_reg_ subsets (Fig. 5B). This suggests that related mechanisms regulate these subsets in DS and highlights another way that IMPACD‘s high-dimensional output can identify organizational principles.

### Advanced immune aging in Down syndrome and type 1 diabetes

Inflammaging describes how immune system remodeling during aging exhibits pro-inflammatory features *(11, 12)*. While people with DS show clinical features of inflammaging, a quantitative understanding of inflammaging in DS remains lacking but is critical to inform mechanistic and translational studies. Our results identified qualitative differences in people with DS suggestive of inflammaging, including decreased naïve T cells, increased CD11c^+^ B cells and increased IL-6 (Fig. 1D-E, 3A and 4E). We sought to use the wide age range of our cohort to investigate how these features change with age in people with and without DS. First, we found that naïve T cells (particularly CD8^+^) decreased faster with age in DS versus TC; memory CD4^+^/CD8^+^ T cells increased with age indistinguishably in both groups (Fig. S4A). Second, CD11c^+^ B cells, which are increased in murine aging and in people with rheumatoid arthritis, are higher in people with DS than TCs, but do not increase with age in either population (Fig. S4B) *(28, 55, 56)*. Third, pro-inflammatory cytokines including TNFα, IL-1β, IL-6, and IL-22 were elevated in serum from people with DS versus TCs as previously reported, but did not increase linearly with age in either cohort (Fig. 4E and S4C-D) *(8)*. These results highlight how individual features of inflammaging can change with age at different rates, which is an important consideration in any modeling endeavor.

Based on these findings, we built three linear models of inflammaging, which we term immune clocks, all trained using only immune subset population data from TC. In our first “unfiltered” model, we leveraged IMPACD’s high-dimensionality output to evaluate each of the 294,061 subsets identified in TC and found 61 informationally non-redundant immune subsets that correlated most linearly with age in TC (p<0.001). We used similarity clustering (p<0.75) to identify 19 representative subsets to prevent overfitting and principal component analysis (PCA) to generate a linear model using these 19 subsets. This “unfiltered” model showed excellent correlation with age (r^2^=0.92), demonstrating the utility of our approach (Fig. 6A). We validated this approach in two ways. First, we generated training/validation datasets from our TC data by withholding data from 5 randomly selected individuals in a validation dataset. We used an identical approach to build a linear model of age with the remaining 23 individuals and interrogated the quality of age prediction in the validation dataset. Over 5 iterations, using different individuals in each validation dataset, we found reproducibly consistent predictions, supporting our approach (Fig. S4E). Second, we used PCA to build a model based on the 27 subsets in our study that were also found to change with age in a previous publication by Alpert et al *(57)*. This “Alpert-filtered” model, generated using an independent set of markers not enriched for linear correlation with age, predicted age with reasonable accuracy (r^2^ = 0.46, Fig. 6A). Lastly, we focused on the 651 subsets identified by IMPACD as differently abundant in DS versus TC. From these, we identified 41 subsets that correlated linearly with age in TC (Fig. 6A). These included subsets of all major cell types with a slight over-representation of CD4^+^ subsets (Fig. S4F). To prevent overfitting, we used similarity clustering (p<0.85) to extract a representative set of 24 subsets (Fig. S4F-G). We used PCA to generate a linear model with these 24 subsets that correlated with age extremely well (“DS-filtered” model, r^2^ = 0.77, Fig. 6A). Therefore, we built three immune clocks that accurately describe how the immune landscape of TC changes with age.

**Fig. 6.**
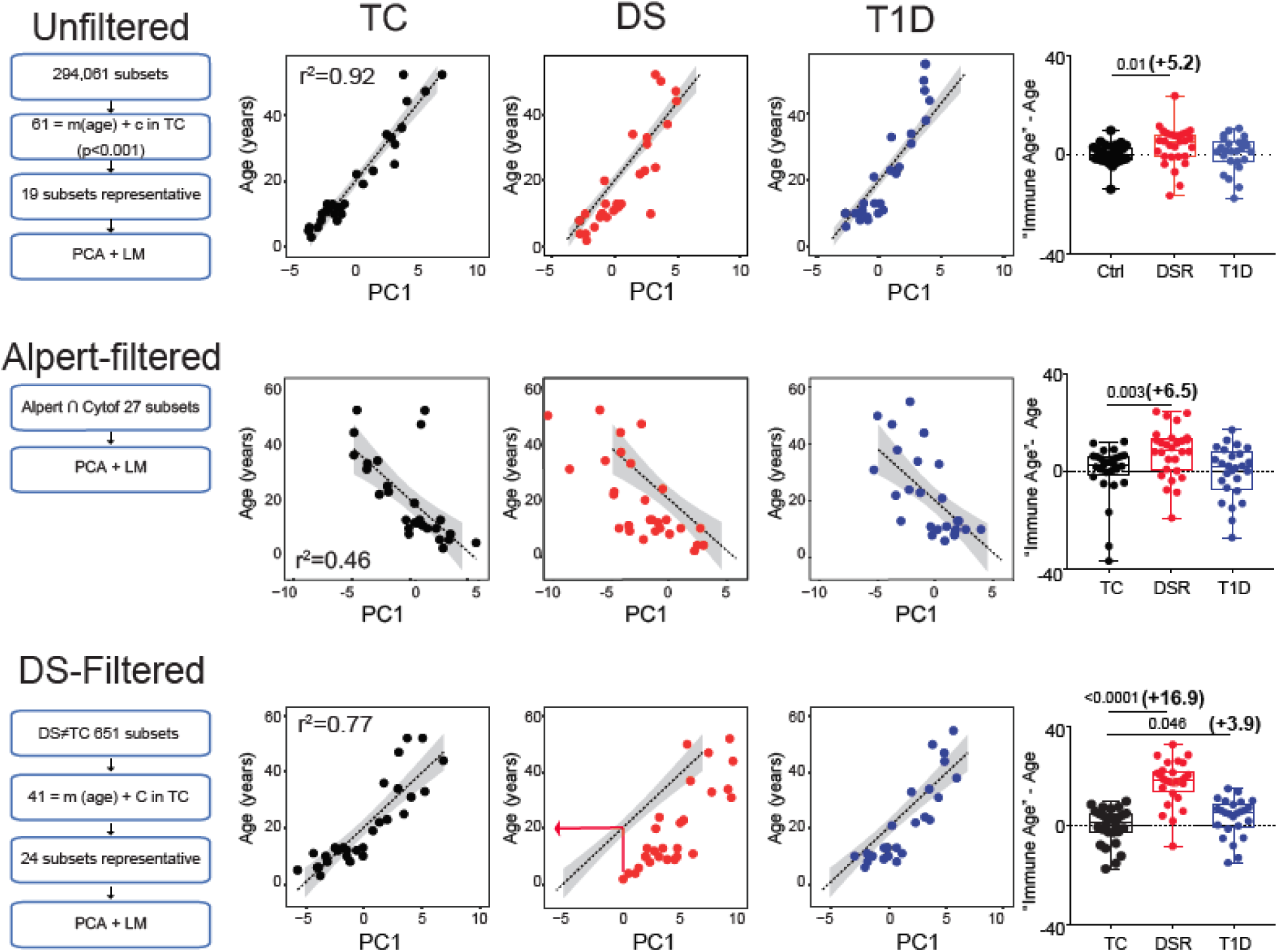
Advanced immune aging in people with Down syndrome. Left, approach to build each linear model of age. All models are trained only on data from controls (TC). Where indicated, subsets that vary linearly with age (i.e. = m(age) + c in TC) are identified and representative subsets identified by similarity clustering as described in the text to prevent overfitting, followed by principal component analysis (PCA) and linear modeling (LM). The TC-derived algorithm is used to predict immune age in people with DS or T1D (middle, red arrow). The difference (predicted immune age – chronological age) is summarized for each cohort (right). n = 28 TC, n = 28 DS, n = 25 T1D, across 5 batches. Mann-Whitney test, p-values shown.

We next used these immune clocks to compare the “immune age” of participants with DS, calculated using their individual immune subset values (red arrow, Fig. 6A), to their actual age. All three immune clocks unanimously showed significantly advanced immune aging in DS (Fig. 6A). The “unfiltered” and “Alpert-filtered” models predicted comparable magnitude of advanced immune aging beginning in childhood (+5.2 and +6.5 years, respectively). This suggests that these models quantitate similar aspects of the immune landscape. Notably, the “DS-filtered” model predicted that the immune system of a person with DS resembles that of a person 16.9 years older on average (Fig. 6A, right). This difference was observed beginning at the earliest age of our cohort, demonstrating again that advanced immune aging in DS begins in childhood (Fig. 6A). This model also predicted linear immune system change with age in DS (r^2^= 0.88, Fig. 6A). The immune age of participants with DS and TC begins to converge past about 35 years of age; whether this reflects a fixed biological endpoint or the upper limit of our training cohort remains to be clearly elucidated. Taken together, these findings exemplify our ability to generate quantitative linear models (immune clocks) that unanimously point to inflammaging (advanced immune aging) in people with DS. Importantly, even though all immune clocks were built on data from the same TCs, comparison of predictions from all three models suggests that the “DS-filtered” model can provide unique DS-relevant biological insight.

To further compare the “unfiltered” and “DS-filtered” models, we assessed PC1 loading coefficients in both models and found that all components contributed equitably, arguing against the dominant effect of few components (Fig. S4H). We next compared features of the subsets that correlated linearly with age in TCs in both models, prior to similarity clustering to ensure an accurate and comprehensive view. We found that root nodes were utilized at similar frequencies in both models (Fig. S4J). Differences were observed in the most commonly used modifier nodes, which were CD161^+^ and Helios^+^ in the “Unfiltered” model and PD-1^+^ and CD73^+^ in the “DS-filtered” model. Terminal nodes used in both models showed some similarities (e.g. high representation of CD38^+^, CD95^+^ and naïve^+^); the “DS-filtered” model showed relative over-representation of naïve^+^, Helios^+^ and CD25^+^ terminal nodes (Fig. S4J). These findings show how selection of different subsets, even within the same major cell types, that vary linearly with age in TC can reveal novel features of aging-related biology.

Finally, we compared the “immune age” of participants with T1D to their actual age. Interestingly, the “DS-filtered” model showed that the immune systems of participants with T1D resemble those of TC subjects who are 3.9 years older on average (Fig. 6A). This difference is greater in younger participants and converges with increasing age as we observed in DS with no consistent bias found in subjects beyond about 30 years of age (Fig. 6A). Neither the “unfiltered” nor the “Alpert-filtered” model showed evidence of advanced immune aging in participants with T1D, although the relative differences in predicted immune age correlated well with the “DS-filtered” model (Fig. 6A and Fig. S4I). Together, these findings point to an association between advanced immune age and T1D and highlight how studying immune dysregulation in DS can highlight novel biology in other autoimmune diseases.

## DISCUSSION

Aging is associated with increased autoimmunity, decreased immune exhaustion, loss of protective immunity and increased pro-inflammatory cytokines, especially IL-6 *(14)*. These features are comprised by the term inflammaging *(12)*. Associated changes in epigenetics, signaling and immune subsets may contribute to autoimmunity risk, although mechanistic details remain incompletely elucidated *(57, 58)*. Drawing causal links to autoimmunity in the general population is further challenged by diverse pathogenic subtypes *(59)*. In this context, Down syndrome represents a unique need and opportunity. People with Down syndrome exhibit clinical, cellular and molecular features of inflammaging, including increased autoimmunity, early-onset Alzheimer’s disease, decreased naïve T cells, altered epigenetics/glycomics and increased serum IL-6 *(2, 6-8, 58, 60)*.Thus, trisomy 21 is a genetic driver of inflammaging in Down syndrome (DS-inflammaging). Specific therapeutic strategies may best mitigate DS-inflammaging-associated diseases in people with Down syndrome and pathophysiologically-related subsets of the general population.

We developed a new software (IMPACD) to interrogate PBMC-CyTOF analysis and built three linear models of immune age (immune clocks) using only typical control data. All three immune clocks quantitatively demonstrate advanced immune aging (i.e. inflammaging) in people with Down syndrome beginning from childhood. Our cohort clarifies that DS-inflammaging is independent of autoimmunity and may help explain why autoimmunity develops more frequently and at younger ages in people with Down syndrome *(3)*. This may also help explain other clinical differences in Down syndrome, including decreased threshold age for poor outcome from SARS-CoV2 infection (40 in Down syndrome versus 60 in the typical population) *(61)*. These findings provide a novel framework to mechanistically investigate how DS-inflammaging impacts other aspects of health in Down syndrome. Responses to vaccination and infection are of particular interest, given that (i) inflammaging was observed even in our youngest participants with Down syndrome, (ii) previous studies suggest impaired vaccine response in Down syndrome and (iii) pulmonary infections represent the leading cause of death in Down syndrome *(62, 63)*. Better understanding DS-inflammaging may guide therapeutic selection and development. Longitudinal studies in larger cohorts will help establish the predictive value of this metric.

Our “DS-filtered” model demonstrates that type 1 diabetes is associated with inflammaging and highlights how studying immune dysregulation in Down syndrome can advance our understanding of other immune-mediated diseases. Dysregulation of B, CD4^+^ T, CD8^+^ T, innate immune and/or islet β cells can drive type 1 diabetes risk and progression *(59)*. Our findings highlight an immunologic commonality between type 1 diabetes and Down syndrome, providing a basis for mechanistic and translational inquiry. Qualitative concordance between all 3 models supports that studying immune aging in type 1 diabetes is likely to be broadly important; how this might stratify prediabetic subjects for risk of progression, prioritize subjects for specific disease-delaying therapy and/or predict response to different therapies is of interest. The similar results of the “unfiltered” and the “Alpert-filtered” models suggests that they reflect similar aspects of aging-related biology. The “DS-filtered” model’s quantitatively distinct results exemplifies how DS-immunodysregulation can inform our understanding of other immune diseases. This use of a signature defined in one context to highlight overlapping biology in a second context is analogous to GSEA analysis of RNAseq data and highlights how IMPACD uniquely allows application of this concept to cytometry data. Utility of this approach should increase exponentially as future experiments generate more reference cytometric signatures.

We found additional interesting features of DS-immunodysregulation. First, our findings suggest that naïve CD4^+^ T cells in Down syndrome are hyperactivated, due in part to elevated IL-6, consistent with data from us and others showing elevated IL-6 in people with Down syndrome *(8)*. Our findings extend the consideration of pathogenic cytokines in Down syndrome beyond type 1 interferons, heretofore the best-known drivers of DS-immunopathology *(9)*. Future work will evaluate how these cytokines coordinately impact naïve CD4^+^ T cell response to activation and Th differentiation. Although the cellular source of increased IL-6 in Down syndrome remains undetermined, IL-6 is linked to autoimmunity in the general population and to early-onset Alzheimer’s disease in Down syndrome, suggesting an immune contribution to DS-Alzheimer’s *(64)*. Second, DS-immunodysregulation centrally impacts NKT cells, including quantitative changes, qualitative remodeling and differential co-regulation with CD4^+^ T cells, suggest overlapping Down syndrome-specific mechanistic pathways. NKT cells are autoimmunity-relevant, although conflicting observations in murine models and in people with type 1 diabetes underscore the need for further study *(65)*. Understanding NKT dysregulation in Down syndrome may inform mechanistic studies in the typical population. Third, B cell compartment remodeling in Down syndrome recapitulates several features of other autoimmune diseases, including (i) increased CD11c^+^ aNAV cells, a major source of antibody secreting cells and serum autoantibodies in SLE *(29)*, (ii) increased CXCR3 and decreased CCR6 in SWM B cells, suggestive of altered migratory potential to inflammation sites in RA *(34)* and SLE *(35)* and (iii) increased PD-1 expression, reminiscent of hyperactive B cells in the joints of patients with RA *(33)*. Whether this reflects quantitative/qualitative defects in T cell help and/or B cell-intrinsic dysregulation remains to be clarified. These findings further emphasize how immune dysregulation in Down syndrome can inform our understanding of other autoimmune diseases.

Our analysis was facilitated by IMPACD, a new software tool we built. IMPACD significantly advances analysis of high-dimensional cytometry data by (i) using digital (manual/automated) gating strategies, (ii) improving the rigor of manual gating analyses, and (iii) performing exhaustive permutational analysis with (iv) zero down-sampling and (v) robust multiple testing correction. Continued interest in innovating digital and manual gating methods is exemplified by approaches including SYLARAS and FAUST *(66, 67)*. Advantages of digital gating methods include longstanding understanding of many markers, ability to adjust for batch effects, and direct translation of results into sorting/analytic strategies for mechanistic studies. IMPACD may be synergistically used with clustering-based algorithms, particularly when digital gating thresholds are less self-evident. Key distinguishing features between IMPACD and clustering approaches include (i) firm control over granularity of the analysis by sequentially evaluating individual markers, (ii) unambiguous definition of cellular subsets for future studies and (iii) zero down-sampling allows robust interrogation of smaller subsets. We show how IMPACD can direct analysis in the context of known biologic frameworks and generate high-density data that can be mined using ‘omics-related methods to highlight novel mechanistic hypotheses. We envision that broader usage will generate more context- and perturbation-specific cellular signatures to enhance functionality while integration with other data modalities will advance hypothesis generation efforts.

## MATERIALS AND METHODS

Detailed methods are included in Supplemental methods.

### Participants and study design

Samples for this study were obtained from participants in the Benaroya Research Institute Immune Mediated Disease Registry. Control participants were selected based on the absence of autoimmune disease or a family history of autoimmunity. The DS cohort was selected to minimize participants with autoimmunity (limited to Hashimoto’s disease in 2 of 28 subjects). Research protocols were approved by the Benaroya Research Institute Institutional Review Board. All participants with DS provided assent. Participants and/or their parents provided written informed consent before participation in the study.

### Statistics

Unless otherwise described, statistical testing (Mann-Whitney, Wilcoxon ranksum testing, linear regression) was performed using Prism (Graphpad).

### Supplementary Materials

Materials and Methods

Fig. S1. Characterization of the cohort.

Fig. S2. IMPACD enables rigorous analysis of manually gated cytometry data.

Fig. S3. Dysregulation of cellular subsets and signaling in T1D and DS.

Fig. S4. Interrogating immune aging in people with DS.

Table S1. CD4-NKT subsets differentially co-regulated in DS.

Table S2. Source of Reagents

## Acknowledgments

We gratefully acknowledge members of the BRI Translational and Diabetes Clinical Research Programs, including Gina Marchesini, Kassidy Benoscek, Marli McCulloch-Olson, Clementine Chalal, Kaytlyn Ly, Laynee Laube, Rebecca Rawlings, Claire Mangan, Catie Wandell, Jenna Snavely, Ezra Graziano, Lisa Miller, McKenzie Lettau, and Jani Klein for subject recruitment and the BRI Clinical core, including Thien-Son Nguyen and Nicole Gilbert for assistance with sample processing and handling. We thank Dr. Hamid Bolouri, Dr. Kirsten Diggins and Tee Bahnson for helpful discussion. We thank Dr. Anne Hocking for editing and proofing the manuscript.

## Author contributions

Conceptualization: BK, JHB, RP

Methodology: BK, JHB, GG, SAL, CS, CJG

Investigation: KL, KGM, AA, GG, KJF, AEW, BK, RP

Visualization: KL, KGM, AA, GG, BK

Funding acquisition: JHB, BK, CJG

Project administration: JHB, BK Supervision: BK, GG

Writing: BK, JHB, KL, KGM, AA, GG, KJF

## Data and materials availability

All data will be made available upon request

