## SupplementalMaterial for "Permutational immune analysis reveals architectural similarities between inflammaging, Down syndrome and autoimmunity"

#### Supplemental Methods

##### Mass Cytometry

Samples were processed by blinded individuals. All 81 samples were analyzed in 5 batches. All members of each age- and sex-matched set were analyzed in the same batch. Each batch included aliquots from a single individual (internal control) to monitor staining variability. Frozen PBMCs were thawed at 37°C in warm complete media (CM, RPMI-HEPES + pyruvate + non-essential amino acids + penicillin-streptomycin + 10% fetal bovine serum). Cells were treated with benzonase nuclease (1:5000 in CM; Sigma) for 5 mins at room temperature (RT). Surface markers: $10^6$ cells were used per stain. Cells were incubated with Cisplatin/ phosphate buffered saline (PBS) solution (1:1000; Enzo Life Sciences) for 1 min at RT. Samples were barcoded by staining with a randomly assigned unique combination of two different Cd-isotope-labelled (of five available) anti-CD45 antibodies. Cells were stained in 100 μL Maxpar Cell staining buffer (CSB, Fluidigm) for 20 mins at RT. Up to 10 (${}^5C_2$) barcoded samples were combined ($3x10^6$ cells per 100 μL) and stained for surface markers for 20 mins at RT. Samples were resuspended in Maxpar Fix and Perm Buffer + DNA Intercalator-Ir (FPBDNAI; Fluidigm, 1:1000, $3x10^6$ cells/100μL) for 1 hour at RT and stored at 4°C until acquisition. Intracellular markers: cells were stimulated with PMA (50ng/mL) + Ionomycin (1μM) in the presence of Brefeldin A (1x) and Monensin (1x) for 3.5 hours at 37°C. Cells were stained with Cisplatin/PBS, barcoded using one Cd-isotope-labelled anti-CD45 antibody per sample, combined (up to 5 = ${}^5C_1$ samples) and stained for surface markers as described above. Cells were fixed using Maxpar Nuclear Antigen Staining Fix (Fluidigm) for 20 mins at RT, stained for intracellular markers in Maxpar Nuclear Antigen Staining Perm (NASP; Fluidigm, $3x10^6$ cells/100 μL NASP) for 30 mins at RT, resuspended in 100 μL FPBDNAI (1:1000, $3x10^6$ cells/100μl) for 1 hour at RT and stored at 4°C until acquisition. Samples were acquired on a Helios mass cytometer (Fluidigm). FACS data was analyzed using FlowJo (TreeStar) and IMPACD (described below).

Absolute cell counts were calculated using contemporaneous clinical complete blood count (CBC) data, available for almost all subjects (26/28 TC, 28/28 DS and 25/25 T1D). The absolute number of PBMCs (#cells/mL of blood) was calculated as sum of the monocyte and lymphocyte counts. Cellular frequencies derived from mass cytometry were then used to calculate #cells/mL of blood of each immune subset.

##### IMPACD (*I*terative *M*achine-assisted *P*ermutational *A*nalysis of *C*ytometry *D*ata)

IMPACD is a software tool, built in the MATLAB environment, designed to enable high-rigor analyses of multiplex immunophenotyping datasets using rapid and deep permutational analyses of digital gating strategies (e.g. manual gating). IMPACD is broadly applicable to immune or non-immune comparisons using mass or flow cytometry. IMPACD analysis is conducted in two stages.

Stage 1: Flowjo is used to manually gate cytometry data and export gating thresholds (XLS file) to IMPACD. IMPACD is compatible with (i) various gate types including histograms, quadrants, rectangles, and polygons, (ii) Boolean operations (e.g. AND, OR and NOT) and (iii) batch-specific threshold definitions. All identified discretization thresholds for each marker and for each batch are reported in an easy-to-read tabulated XLS file for review. This allows for highly intentional retention of only selected variation, such as batch effects (i.e. highest-rigor manual gating analysis). Using finalized threshold values, sample FCS files are automatically read and each cell is individually digitized in a high dimensional multiplexed discrete space according to its expression of each marker. IMPACD is compatible with both binary (positive or negative) and higher-order (e.g. low, intermediate, high) definitions of marker expression.

Stage 2: IMPACD performs exhaustive permutational analysis of all markers without any down-sampling to elucidate cellular subsets with statistically significant frequency differences between cohorts, including robust multiple test correction. A "root path" is defined for each cell type of interest as a sequence of expressed markers (e.g. $CD4^+$ NKTs as $/CD14^-/CD3^+/CD19^-$ $/CD56^{hi+}/CD4^+$/CD8- cells). Descendant subsets are then defined by sequential addition of remaining markers/gates, and described by a path including "root", "modifier(s)" and "terminal" nodes. The number of cells in each subset in each subject are enumerated. Subsets <10 cells (median) are excluded to minimize noise. Significance of difference is assessed using the nonparametric Wilcoxon ranksum test and p-values are adjusted for multiple hypothesis testing by Benjamini-Hochberg method. Significantly scoring subsets with a single modifier node are further cross-examined against the subset with the same root path and terminal node but no modifier node (e.g if /A+/B+/C+ scores, compare against /A+/C+). Likewise, significantly scoring subsets with two modifier nodes are compared against the subsets with the same root path, terminal node and one of the two modifier nodes (e.g. if /A+/B+/C+/D+ scores, compare against /A+/B+/D+ and /A+/C+/D+). Only subsets with significant information gain (as assessed by Wilcoxon ranksum-based statistically significant difference in cell frequencies) are reported.

**Polyexpression analysis**

IMPACD digitizes each cell in a multidimensional discrete space, allowing us to exhaustively test for statistically significant differences in all combinations of marker expression in a computationally inexpensive manner. Naïve $CD4^+$ T cells were analyzed for polyexpression of CXCR3, CD39, CD73, CD38, CD62L, ICOS, and TIGIT. Using these seven markers, 128 different combinations of marker expression were enumerated in TC, DS and T1D. Cell frequencies were expressed as percentage of naïve $CD4^+$ T cells. Significance of difference in frequencies between two cohorts was assessed using the nonparametric Wilcoxon ranksum test and p-values were adjusted for multiple hypothesis testing by Benjamini-Hochberg method.

**Clustering analyses**

Naïve $CD4^+$ T cells were gated manually in FlowJo and exported FCS files were analyzed using the Cytofkit and flowCore packages in R *(68)*. Clustering markers used were: CXCR3, ICOS, TIGIT, CD62L, CD73, CD38 and CD39. Marker intensity values were arcsinh transformed with cofactor = 5 and events were subsampled to 1,000 events per sample. Clustering was performed using the FlowSOM function from the flowCore package without scaling. Repeat consensus clustering identified 12 clusters as the modal number; 6 small clusters (<1% of cells in TC and <2.5% of cells in DS/T1D) were excluded and the 6 remaining clusters were used in downstream analyses. The proportions of cells from each sample falling into each cluster were statistically compared using quasibinomial logistic models with age as a covariate in the model. Resulting p-values were corrected for multiple comparisons using the Holm correction. Thresholds for CD62L (>4.7) and CD38 (>9.9) positivity were as defined by manual gating.

**Network analysis**

Differentially co-expressed modules were identified using the diffcoexp package in R *(69)*. From the 651 subsets identified by IMPACD as different between people with and without DS, this algorithm (i) separately calculated correlation between each pair of subsets in participants with and without DS, (ii) compared all pairs found to be significantly correlated in either cohort and (iii) reported pairs found to be differentially correlated by Fisher's Z-transformation. As a control, we performed a random simulation experiment, where we randomly reassigned the cohort label of the 28 control and 28 DS subjects and identified differentially correlated pairs over 10,000 iterations. This allowed us to calculate FDR for each category of pairs as # instances pairs in random simulation ≥ # pairs in actual data. The network model was generated using Cytoscape *(70)*.

**Immune age modeling**

Subset selection, "unfiltered" model: All 294,061 permutations of marker expression (subsets) found by IMPACD (>10 cells/subset) in TC were interrogated. Data for each subset for the control cohort was tested for linear correlation with age by mixed linear model in R. Sex and batch number were included as covariates in the linear model to generate FDR-adjusted p-values. The subsets that were most significantly correlated with age ($p<0.001$) were selected. To minimize informational redundancy, selection began with the lowest p-value and each unique root + terminal node combination was selected only once, identifying 61 subsets. Pairwise correlation clustering was performed for all markers, using the rdist package in R to calculate distance via distance-sampling. Hierarchical clustering ($r<0.75$) was then used divide these 61 subsets into 19 clusters. We selected from each of these 19 clusters the subset that showed highest correlation with age. This produced a smaller set of 19 representative subsets and prevented overfitting in our linear model.

Subset selection, "DS-filtered" model: Beginning with the 651 subsets identified by IMPACD as different between people with and without DS, we tested data from TC as described above for linear correlation with age ($p<0.05$). The 41 subsets identified were analyzed by hierarchical clustering ($r<0.9$) to identify 24 representative subsets.

Linear modeling vs age: For each subset, CyTOF data for control participants was min-max normalized and z-scale normalized. PCA was performed in R. PC1 was the most correlated with age in all models. PC1 and age were used to generate a linear model using only data from TC participants. Data from DS/T1D participants was normalized against the control participants and projected onto the PCs defined using control participant data. "Immune age" of each participant was calculated using the participant's PC1 value and the linear model generated using control data.

Training set validation: The control cohort was split into 5 unique training/validation sets. Each validation set comprised 5 unique individuals, 3 <14 years old and 2 >18 years old. Data from non-validation individuals was used for training. Linear modeling was performed as described using the training dataset and immune age of the validation set calculated.

**Phospho-flow Assay**

Per condition tested, $4x10^5$ thawed PBMCs were stained for viability (Zombie NIR, Biolegend) and selected surface markers (CXCR3, CCR6, CXCR5, CD62L) for 20 min at RT in PBS and stimulated with cytokine (250 IU/mL IFNα, PBL Assay Science or 1 ng/ml IL-6, BD Biosciences) or no-cytokine control in serum-free media (X-VIVO15, Lonza) for 30 min at 37°C. Cells were fixed using Fix buffer I (BD Bioscience) for 15 mins at 37°C, permeabilized with ice-cold Perm buffer III (BD Bioscience) for 30 mins on ice and stained for remaining markers (CD3, CD4, CD8, CD56, CD27, CD45RA, CD25, pSTAT1, pSTAT3, pSTAT5) in FACS buffer for 45 mins at RT. Samples were acquired on an LSRII (BD Biosciences).

**Cytokine stimulation of T cells**

$CD3^+$ cells were isolated from thawed TC PBMCs by negative magnetic separation (Human Pan T Cell Isolation Kit, Miltenyi). $6x10^5$ $CD3^+$ cells were plated per condition in XVIVO15 media (Lonza) ± cytokine (500 IU/mL IFNα or 1 ng/ml IL-6) and incubated at 37°C for 6 days. Cells were stained for surface markers (CD45RA, CD95, CD38, CXCR3, CD62L, CD27, CD73, CD39, PD-1, Zombie/NIR) in PBS for 20 mins at RT, fixed with BD Cytofix/Cytoperm for 5 min at RT, stained for CD4 and CD8 in Perm Buffer (1X, Invitrogen) for 20 mins at RT and acquired on an LSRII.

**Mesoscale Discovery**

Serum was thawed at 37°C and cytokine concentration was measured using V-PLEX Th17 PANEL 1 Human Kit and V-PLEX Proinflammatory Panel 1 Human Kit (MSD #K15085D-1 and K15049D-1) per manufacturer's instructions.

**CMV ELISA**

Serum was thawed at 37°C and CMV IgG was measured using Human Anti-Cytomegalovirus IgG ELISA Kit (Abcam #ab108639) per manufacturer's instructions.

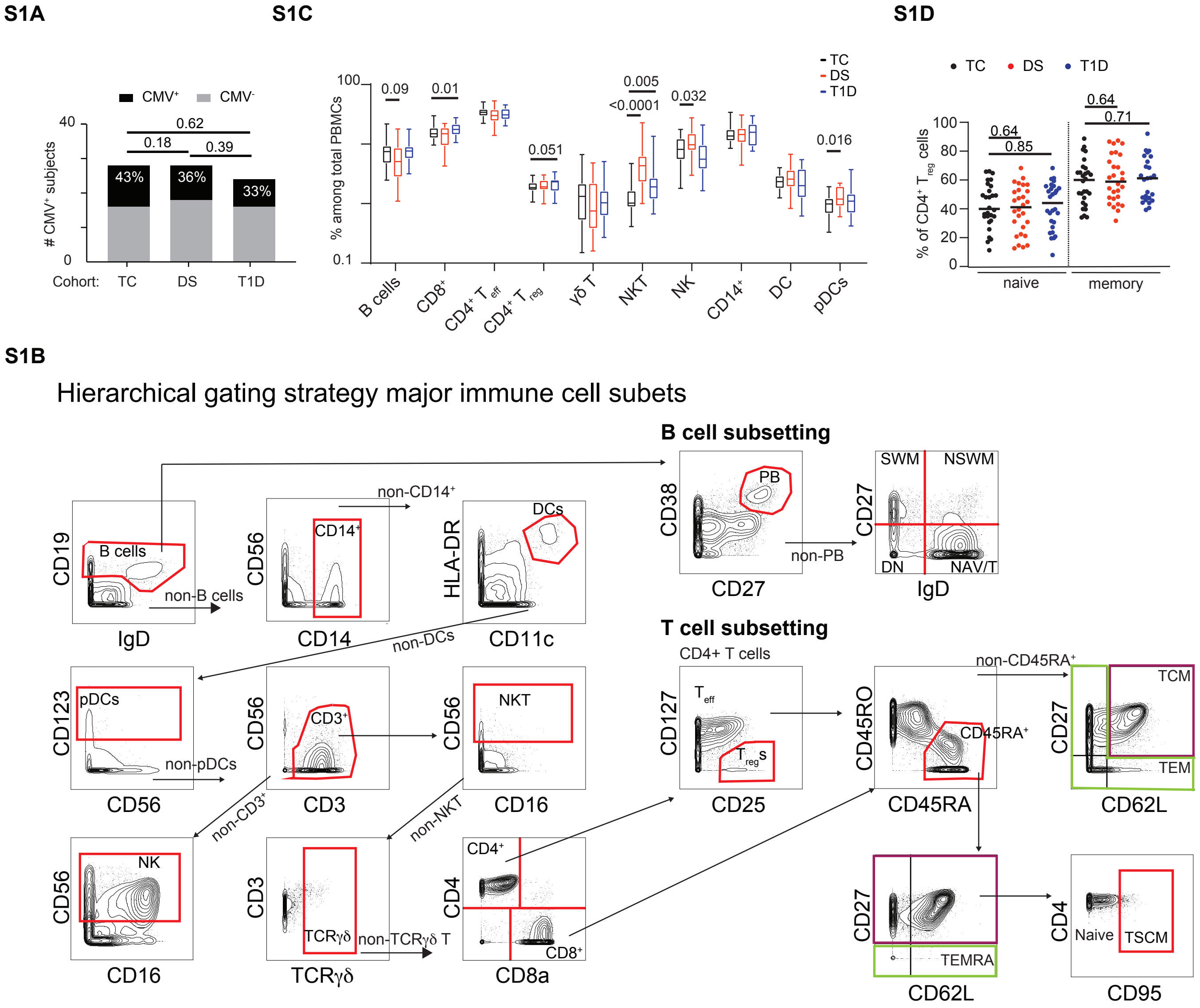


**Fig. S1. Characterization of the cohort.** (**A**) Number and percent of subjects in each cohort who were CMV seropositive. (**B**) Hierarchical gating strategy for major immune subsets. (C) Frequencies of major cell types amongst total PBMCs in each cohort. (D) Frequency of naïve and memory cells amongst Tregs. n = 28 TC, n = 28 DS, n = 25 T1D, across 5 batches. Mann-Whitney test, p-values shown.

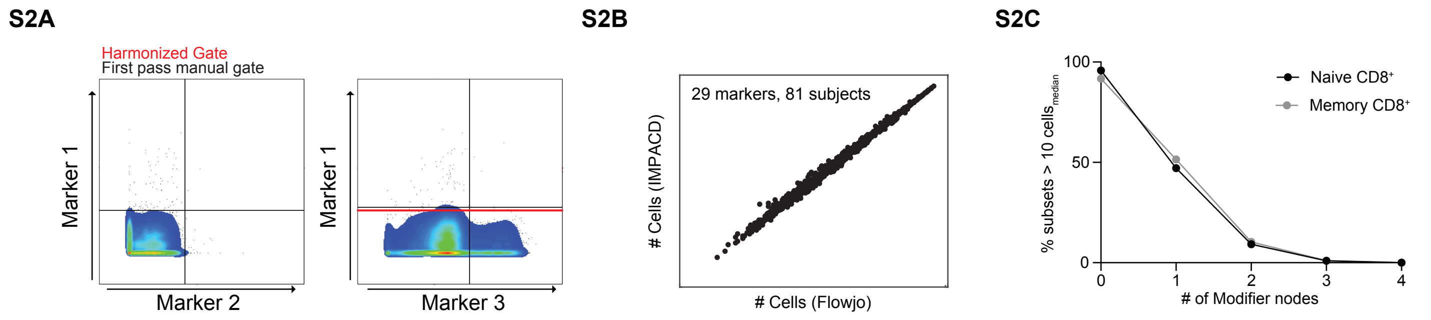


**Fig. S2. IMPACD enables rigorous analysis of manually gated cytometry data.** (**A**) IMPACD reveals when different thresholds are used for one marker in manual gating analysis, as exemplified by the black line in the right plot. This allows for harmonization of inadvertent differences (red line). (**B**) Comparing the number of cells positive for each of 29 markers in all 81 subjects, as determined by Flowjo versus IMPACD, shows comparable results. (**C**) Frequency of subsets with median >10 cells with increasing number of modifier nodes.

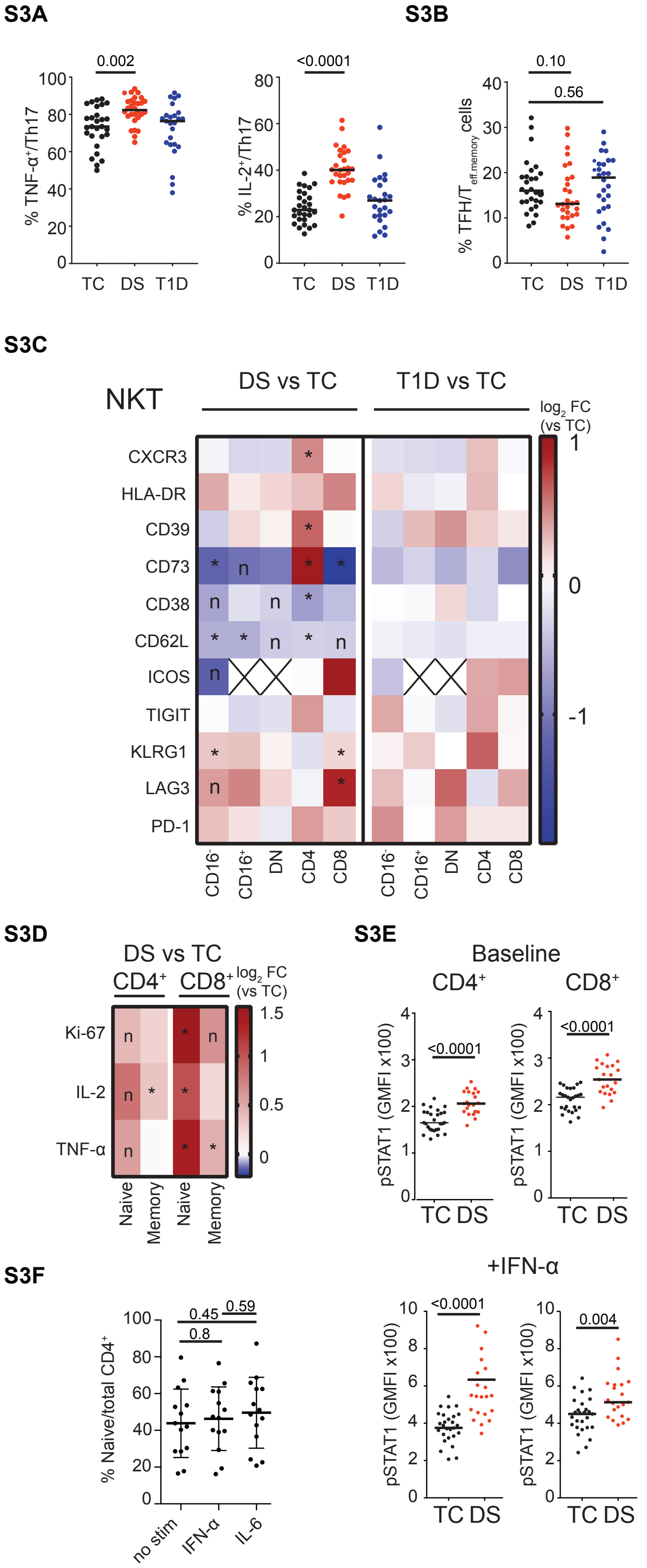


**Fig S3. Dysregulation of cellular subsets and signaling in T1D and DS.** (**A**) Frequency of TNF + and IL-2+ cells amongst Th17 (IL-17 +CD4+ memory T) cells. (**B**) Frequency of (CXCR5+) TFH cells amongst CD45RO+CD4+ Teff (Teff.memory) cells. (**C**) Abundance of each NKT cell subset, in either DS or T1D versus TC, expressing each of several selected markers. (**D**) Abundance of each T cell subset, expressing Ki67, IL-2 or TNF , in DS versus TC. (**C-D**) Heatmap shows the ratio of median percentage of marker-expressing cells. Wilcoxon ranksum test with Benjamini-Hochberg correction, *, FDR<0.05; n, nominal p<0.05; cross, median #cells<10. (**E**) Baseline and IFN -induced pSTAT1 is increased in naïve CD4+ and CD8+ T cells from people with DS. (**F**) Effect of IFN or IL-6 treatment on naïve CD4+ T cell frequency. n = 14 TC. (A-E) n = 28 TC, n = 28 DS, n = 25 T1D, across 5 batches. (**A, B, E-F**) Mann-Whitney test, p-values shown.

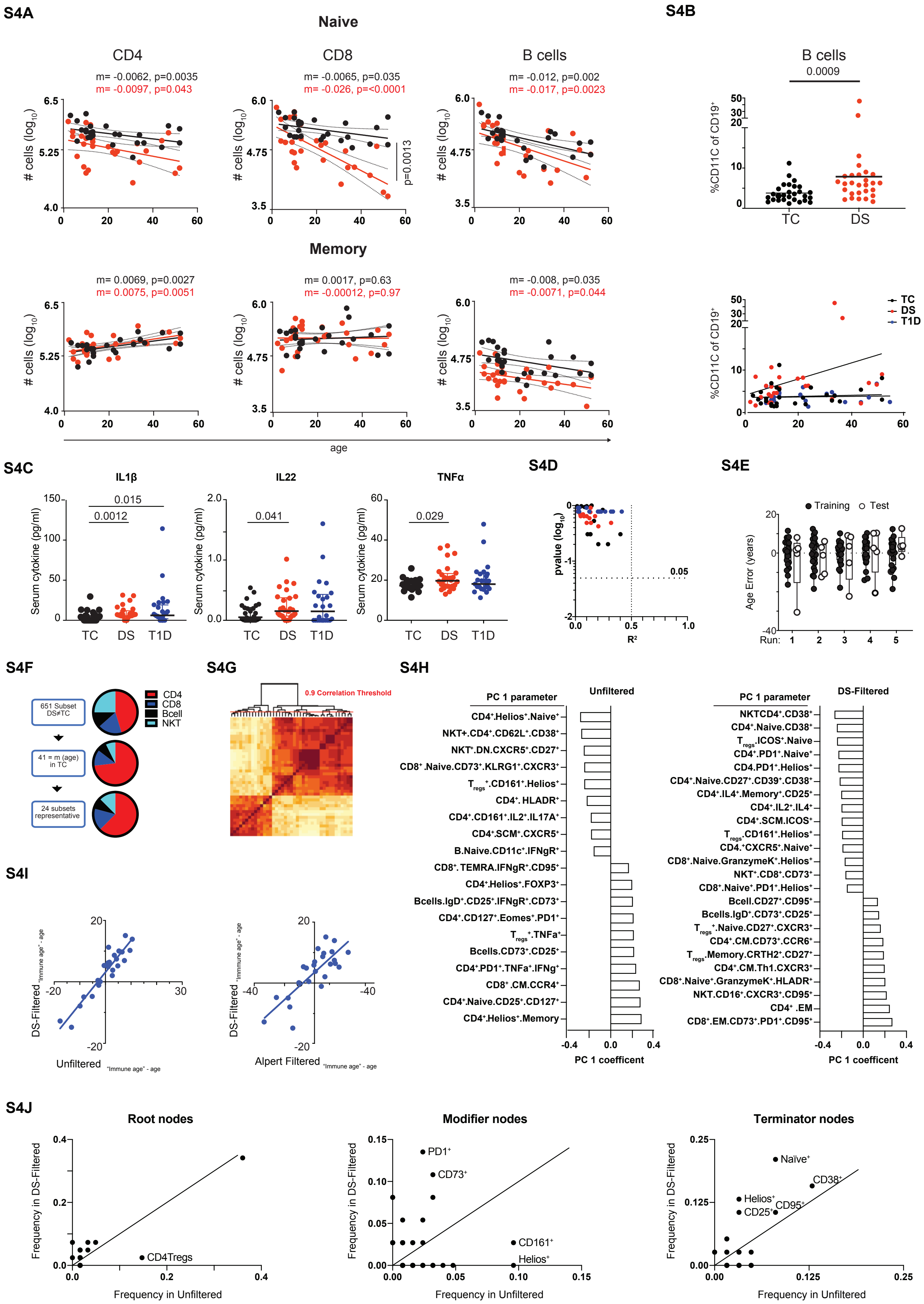

S4A
Naive
CD4
CD8
B cells
m= -0.0062, p=0.0035
m= -0.0097, p=0.043
m= -0.0065, p=0.035
m= -0.026, p=<0.0001
m= -0.012, p=0.002
m= -0.017, p=0.0023
p=0.0013
### cells (log10)
Memory
m= 0.0069, p=0.0027
m= 0.0075, p=0.0051
m= 0.0017, p=0.63
m= -0.00012, p=0.97
m= -0.008, p=0.035
m= -0.0071, p=0.044
age
S4B
B cells
0.0009
%CD11C of CD19+
TC
DS
T1D
S4C
IL1β
IL22
TNFα
0.015
0.0012
0.041
0.029
Serum cytokine (pg/ml)
S4D
pvalue (log10)
R²
0.05
S4E
Training
Test
Age Error (years)
Run:
S4F
651 Subset DS≠TC
41 = m (age) in TC
24 subsets representative
CD4
CD8
Bcell
NKT
S4G
0.9 Correlation Threshold
S4H
PC 1 parameter
Unfiltered
DS-Filtered
CD4+.Helios+.Naive+
NKT+.CD4+.CD62L+.CD38+
NKT+.DN.CXCR5+.CD27+
CD8+.Naive.CD73+.KLRG1+.CXCR3+
Tregs+.CD161+.Helios+
CD4+.HLADR+
CD4+.CD161+.IL2+.IL17A+
CD4+.SCM+.CXCR5+
B.Naive.CD11c+.IFNgR+
CD8+.TEMRA.IFNgR+.CD95+
CD4+.Helios+.FOXP3+
Bcells.IgD+.CD25+.IFNgR+.CD73+
CD4+.CD127+.Eomes+.PD1+
Tregs+.TNFa+
Bcells.CD73+.CD25+
CD4+.PD1+.TNFa+.IFNg+
CD8+.CM.CCR4+
CD4+.Naive.CD25+.CD127+
CD4+.Helios+.Memory
NKTCD4+.CD38+
CD4+.Naive.CD38+
Tregs.ICOS+.Naive
CD4+.PD1+.Naive+
CD4.PD1+.Helios+
CD4+.Naive.CD27+.CD39+.CD38+
CD4+.IL4+.Memory+.CD25+
CD4+.IL2+.IL4+
CD4+.SCM.ICOS+
Tregs.CD161+.Helios+
CD4.+CXCR5+.Naive+
CD8+.Naive.GranzymeK+.Helios+
NKT+.CD8+.CD73+
CD8+.Naive+.PD1+.Helios+
Bcell.CD27+.CD95+
Bcells.IgD+.CD73+.CD25+
Tregs+.Naive.CD27+.CXCR3+
CD4+.CM.CD73+.CCR6+
Tregs.Memory.CRTH2+.CD27+
CD4+.CM.Th1.CXCR3+
CD8+.Naive+.GranzymeK+.HLADR+
NKT.CD16+.CXCR3+.CD95+
CD4+ .EM
CD8+.EM.CD73+.PD1+.CD95+
PC 1 coefficent
S4I
DS-Filtered "Immune age" - age
Unfiltered "Immune age" - age
Alpert Filtered "Immune age" - age
S4J
Root nodes
Modifier nodes
Terminator nodes
Frequency in DS-Filtered
Frequency in Unfiltered
CD4Tregs
PD1+
CD73+
CD161+
Helios+
Naive+
CD38+
Helios+
CD25+
CD95+

**Fig. S4. Interrogating immune aging on people with DS. (A)** Evaluating how absolute numbers of naïve and memory B/T subsets change with age in people with DS (red) and controls (black). Linear regression analysis showing rate of change (gradient, m) together with F-test significance of correlation. (**B**) CD11c+ B cells are increased in people with DS. (**C**) Concentration of selected pro-inflammatory cytokines in serum from people with DS or T1D versus controls (TC). (D) Volcano plot summarizing linear correlation of all cytokines measured with age in people with DS and controls. Dotted lines denote cutoffs of $p<0.05$ and $r2>0.5$. E. Comparing precision, in training and validation subsets, of 5 independent iterations to generate a predictive model of age in TC using immune subsets. F. Proportion of major subsets in each stage of "DS-filtered" linear model development. G. Visual representation of correlation clustering to reduce the "DS-filtered" model's 41 subsets to 24 representative subsets. H. Comparable loading coefficients for all components of both the "unfiltered" and "DS-filtered" models. I. Correlation between immune age advancement calculated using the "unfiltered", "Alpert-filtered" and "DS-filtered" models. J. Comparing frequency of individual root, modifier and terminal nodes in the source 61 and 41 subsets (that correlate linearly with age in TC) for the Unfiltered and DS-filtered models respectively. n = 26 (A) or n = 28 (B-I) TC, n = 28 DS, n = 25 T1D, across 5 batches. (B-C) Mann-Whitney test, p-values shown.

Table S1.CD4-NKTsubsets differentially regulated in DS.

| Subset 1 | Subset 2 | Correlation within control cohort | Correlation within Down syndrome cohort | FDR adjusted nominal p diffcoex (fishers z-transformation) | cor.diff | Nominal p control cohort (pearson corr) | Nominal p control Down syndrome (pearson corr) | Nominal p diffcoex (fishers z-transformation) | FDR adjusted nominal p control cohort (pearson corr) | FDR adjusted nominal p control Down syndrome (pearson corr) |
|---|---|---|---|---|---|---|---|---|---|---|
| CD4TcellsTeff..Tregsneg.PD1plus.Eomesplus | CD4TcellsTeffCRTH2plus | 0.45287224 | -0.601550657 | 0.076153091 | -1.054422898 | 0.015521008 | 0.000709099 | 2.84E-05 | 0.172553059 | 0.01936839 |
| CD4TcellsTregs_Memory_CCR6plus.CD103plus.CD27plus | CD4TcellsTeff_EM_ICOSplus.CRTH2plus | -0.155372044 | -0.877545413 | 0.062282975 | -0.722173368 | 0.429831543 | 8.77E-10 | 1.94E-05 | 0.774304203 | 2.59E-07 |
| CD4TcellsTregs_Memory_CD103plus.CD39plus.CD27plus | CD4TcellsTeff_EM_ICOSplus.CRTH2plus | -0.005907781 | -0.903978037 | 0.006428154 | -0.898070256 | 0.976197882 | 4.34E-11 | 1.44E-07 | 0.993837076 | 1.95E-08 |
| CD4TcellsTeff.Tregsneg.PD1plus.Eomesplus | CD4TcellsTeff_EM_Th17_DS.CXCR3neg.CCR6plus.CRTH2plus.CD27plus | 0.589847487 | -0.531591787 | 0.032283081 | -1.121439274 | 0.000954822 | 0.003600696 | 7.14E-06 | 0.030673626 | 0.062794835 |
| CD4TcellsTeffTregsneg.PD1plus.Naiveplus | CD4TcellsTeff_EM_Th2_DS.CXCR3neg.CCR6neg.CCR4plus.PD1plus | 0.691274426 | -0.293551055 | 0.087366396 | -0.984825481 | 4.64E-05 | 0.129485268 | 4.58E-05 | 0.003591053 | 0.483538122 |
| CD4TcellsTeff_CM_CD73plus.CCR6plus | CD4TcellsTeff_Naive_CXCR5plus.TIGITplus | -0.432396444 | 0.599451236 | 0.085923417 | 1.031847681 | 0.021561002 | 0.000748603 | 4.43E-05 | 0.206762768 | 0.020153416 |
| CD4TcellsTregs_Heliosneg.CD161plus | CD4TcellsTeff_CXCR5plus.TIGITplus | -0.260651817 | 0.703390068 | 0.089514915 | 0.964041885 | 0.180358849 | 2.98E-05 | 5.50E-05 | 0.560673281 | 0.001633476 |
| CD4TcellsTeff_Memory_Tregsneg.Memoryplus | CD4TcellsTregs_Heliosneg.CD161plus | 0.88024211 | 0.018078025 | 0.014798737 | -0.862164084 | 6.67E-10 | 0.927249606 | 1.56E-06 | 3.64E-07 | 0.980374971 |
| CD4TcellsTeff.Tregsneg.IL4plus.Memoryplus.CD25plus | CD4TcellsTregs_Memory_CCR6plus.CD103plus.CD27plus | 0.478933474 | -0.540811773 | 0.097631326 | -1.019745247 | 0.009925122 | 0.002964313 | 6.77E-05 | 0.133980008 | 0.05502376 |
| CD4TcellsTeff_Memory_Tregsneg.Memoryplus.IL4plus.CD25plu | CD4TcellsTregs_Memory_CCR6plus.CD103plus.CD27plus | 0.478933474 | -0.540811773 | 0.097631326 | -1.019745247 | 0.009925122 | 0.002964313 | 6.77E-05 | 0.133980008 | 0.05502376 |
| CD4TcellsTeff.Tregsneg.PD1plus.Eomesplus | CD4TcellsTregs_Memory_CRTH2plus.CD27plus | 0.616330983 | -0.561746039 | 0.014798737 | -1.178077022 | 0.000478887 | 0.00186669 | 1.68E-06 | 0.01942121 | 0.039759756 |
| CD4TcellsTeff_Tregsneg.PD1plus.Granzyme_Kplus | CD4TcellsTregs_Memory_CRTH2plus.CD27plus | 0.66989805 | -0.359144748 | 0.076039343 | -1.029042798 | 9.64E-05 | 0.060525774 | 2.73E-05 | 0.006232064 | 0.337839558 |
| CD8Tcells_Naive_Granzyme_Bplus.Eomesneg.Tbetplus | CD4TcellsTeff_EM_Th17_CXCR3neg.CCR6plus.HLADRplus.CD161neg.CD103plus | 0.264763294 | -0.784238634 | 0.020297806 | -1.049001928 | 0.173335157 | 7.89E-07 | 2.69E-06 | 0.551355122 | 8.18E-05 |
| CD8Tcells_Naive_Granzyme_Bplus.Heliosneg.Tbetplus | CD4TcellsTeff_EM_Th17_CXCR3neg.CCR6plus.HLADRplus.CD161neg.CD103plus | 0.222560954 | -0.767908299 | 0.046600575 | -0.990469253 | 0.254979403 | 1.84E-06 | 1.14E-05 | 0.641540011 | 0.000165443 |
| CD8Tcells_Naive.Granzyme_Bplus.IFNgneg.Tbetplus | CD4TcellsTeff_EM_Th17_CXCR3neg.CCR6plus.HLADRplus.CD161neg.CD103plus | 0.304209429 | -0.761512224 | 0.022542099 | -1.065721653 | 0.115510606 | 2.52E-06 | 3.39E-06 | 0.464366726 | 0.00021558 |
| CD8Tcells_Naive_Granzyme_Bplus.Tbetplus | CD4TcellsTeff_EM_Th17_CXCR3neg.CCR6plus.HLADRplus.CD161neg.CD103plus | 0.241151478 | -0.759505256 | 0.046600575 | -1.000656734 | 0.21637707 | 2.78E-06 | 1.15E-05 | 0.60307437 | 0.000233264 |
| CD8Tcells_EM_TEMplus.CD73plus.PD1plus.CD95plus | CD4TcellsTregs_Memory_CRTH2plus.CD27plus | 0.651715306 | -0.401648331 | 0.063936462 | -1.053363637 | 0.000171979 | 0.034127482 | 2.08E-05 | 0.009531245 | 0.251796691 |
| CD4TcellsTeff_CM_nonTh_CXCR3neg.CCR6neg.CCR4neg.TIGITp | Bcells_Plasmablasts_CXCR5plus.CD95plus | 0.356370038 | -0.651439409 | 0.087970042 | -1.007809447 | 0.062687166 | 0.000173445 | 4.75E-05 | 0.354408173 | 0.006520359 |
| CD4TcellsTeff_Naive.CXCR3plus | BcellsnonPB_DblNeg_DS.nonPBplus.CD27neg.IgDneg.CCR6plus | -0.571755723 | 0.450300028 | 0.093738387 | 1.022055752 | 0.00148021 | 0.016192422 | 5.98E-05 | 0.040975453 | 0.164865815 |
| CD4TcellsTregs_Memory_CD73plus.PD1plus | BcellsnonPB_SwitchedMemory_CD27plus.IgDneg.CD95plus | -0.641169532 | 0.370709996 | 0.087970042 | 1.011879528 | 0.00023649 | 0.052135784 | 4.83E-05 | 0.011921724 | 0.313694045 |
| NKTcells_CD8SP_PD1plus.CXCR3neg.PDL1plus | CD4TcellsTeff_Tregsneg.Naive.IL17Aplus | -0.165488009 | 0.778321068 | 0.062282975 | 0.943809077 | 0.400022344 | 1.08E-06 | 1.94E-05 | 0.755377198 | 0.00010646 |
| NKTcells_CD8SP_PD1plus.CXCR3neg.PDL1plus | CD4TcellsTeff_Tregsneg.Naive.IL17Aplus | -0.211007132 | 0.760195378 | 0.062282975 | 0.97120251 | 0.28110842 | 2.69E-06 | 1.86E-05 | 0.665691194 | 0.000227029 |
| NKTcells_CD8SP_CD161neg.CXCR3neg.CD73plus | CD4TcellsTeff_EM_Th17_CXCR3neg.CCR6plus.HLADRplus.CD103plus | -0.280390052 | 0.684159199 | 0.098660571 | 0.964549251 | 0.148397101 | 5.96E-05 | 6.97E-05 | 0.516286773 | 0.002824304 |
| NKTcells_CD8SP.CD38plus.CXCR3neg.CD73plus | CD4TcellsTeff_EM_Th17_CXCR3neg.CCR6plus.HLADRplus.CD103plus | -0.350614317 | 0.730775572 | 0.026985013 | 1.081389889 | 0.067361731 | 1.01E-05 | 4.56E-06 | 0.366259833 | 0.000659488 |
| NKTcells_CD8SP.CCR6neg.CXCR3neg.CD73plus | CD4TcellsTeff_EM_Th17_CXCR3neg.CCR6plus.HLADRplus.CD161neg.CD103plus | -0.398626642 | 0.62797962 | 0.085193324 | 1.026606262 | 0.035627807 | 0.000346564 | 4.10E-05 | 0.268225931 | 0.011155373 |
| NKTcells_CD8SP_CD161neg.CXCR3neg.CD73plus | CD4TcellsTeff_EM_Th17_CXCR3neg.CCR6plus.HLADRplus.CD161neg.CD103plus | -0.390329021 | 0.667629973 | 0.057945275 | 1.057958994 | 0.040021337 | 0.000103874 | 1.64E-05 | 0.283897079 | 0.004373562 |
| NKTcells_CD8SP.CD38plus.CCR4neg.CD73plus | CD4TcellsTeff_EM_Th17_CXCR3neg.CCR6plus.HLADRplus.CD161neg.CD103plus | -0.375878699 | 0.677413807 | 0.057945275 | 1.053292506 | 0.048695495 | 7.51E-05 | 1.62E-05 | 0.313945493 | 0.003386502 |
| NKTcells_CD8SP.CD38plus.CD62Lplus.CD73plus | CD4TcellsTeff_EM_Th17_CXCR3neg.CCR6plus.HLADRplus.CD161neg.CD103plus | -0.369196422 | 0.665874653 | 0.07228028 | 1.035071075 | 0.053178583 | 0.000109974 | 2.55E-05 | 0.327826327 | 0.004577045 |
| NKTcells_CD8SP_CD38plus.CD73plus | CD4TcellsTeff_EM_Th17_CXCR3neg.CCR6plus.HLADRplus.CD161neg.CD103plus | -0.387048615 | 0.645845471 | 0.079416529 | 1.032894086 | 0.041873174 | 0.000205643 | 3.19E-05 | 0.290922024 | 0.007451429 |
| NKTcells_CD8SP_CD38plus.CXCR3neg.CD73plus | CD4TcellsTeff_EM_Th17_CXCR3neg.CCR6plus.HLADRplus.CD161neg.CD103plus | -0.470302765 | 0.699091313 | 0.014257502 | 1.169394078 | 0.011552091 | 3.49E-05 | 1.15E-06 | 0.145992497 | 0.001843431 |
| NKTcells_CD8SP_CD38plus.TIGITneg.CD73plus | CD4TcellsTeff_EM_Th17_CXCR3neg.CCR6plus.HLADRplus.CD161neg.CD103plus | -0.326167815 | 0.673942487 | 0.085728628 | 1.000110302 | 0.090282086 | 8.43E-05 | 4.34E-05 | 0.416888898 | 0.003716034 |
| NKTcells_CD8SP.CD62Lplus.CXCR3neg.CD73plus | CD4TcellsTeff_EM_Th17_CXCR3neg.CCR6plus.HLADRplus.CD161neg.CD103plus | -0.36985081 | 0.75262865 | 0.014798737 | 1.12247946 | 0.052725739 | 3.84E-06 | 1.34E-06 | 0.326258183 | 0.000301887 |
| NKTcells_CD16neg.CD62Lplus.IFNgRplus.KLRG1plus | CD4TcellsTeff_EM_Th2_CXCR3neg.CCR6neg.CCR4plus.PDL1plus.CD27plus | -0.517320731 | 0.707560425 | 0.006741308 | 1.224881156 | 0.004814836 | 2.54E-05 | 2.69E-07 | 0.086994816 | 0.001431867 |
| NKTcells_CD16neg.CD95plus.ICOSplus | CD4TcellsTeff_EM_Th2_CXCR3neg.CCR6neg.CCR4plus.PDL1plus.CD27plus | 0.387337322 | -0.771686655 | 0.007775514 | -1.159023977 | 0.041707507 | 1.52E-06 | 4.04E-07 | 0.290281448 | 0.000140858 |
| NKTcells_CD16neg.IFNgRplus.CCR6neg.KLRG1plus | CD4TcellsTeff_EM_Th2_CXCR3neg.CCR6neg.CCR4plus.PDL1plus.CD27plus | -0.460638389 | 0.622459991 | 0.055168716 | 1.08309838 | 0.01363214 | 0.000404602 | 1.43E-05 | 0.160183346 | 0.012573977 |
| NKTcells_CD8SP_CD62Lplus.IFNgRplus.KLRG1plus | CD4TcellsTeff_EM_Th2_CXCR3neg.CCR6neg.CCR4plus.PDL1plus.CD27plus | -0.355259464 | 0.770392738 | 0.0137936 | 1.125652202 | 0.063568893 | 1.63E-06 | 8.48E-07 | 0.356794454 | 0.000148891 |
| NKTcells_CD8SP_IFNgRplus.CCR6neg.KLRG1plus | CD4TcellsTeff_EM_Th2_CXCR3neg.CCR6neg.CCR4plus.PDL1plus.CD27plus | -0.307381854 | 0.705091777 | 0.070218099 | 1.012473631 | 0.111575106 | 2.79E-05 | 2.39E-05 | 0.457525836 | 0.001548774 |
| NKTcells_CD16neg.CD161neg.PD1plus | CD4TcellsTeff_Memory_Tregsneg.IL17Aplus.IL2plus | 0.416605272 | -0.594547692 | 0.097631326 | -1.011152964 | 0.027431316 | 0.000848435 | 6.64E-05 | 0.234917354 | 0.022092359 |
| NKTcells_CD16neg_PD1plus.CXCR3neg.CCR6plus | CD4TcellsTeff_Naive_TIGITplus | -0.367240409 | 0.749859414 | 0.014798737 | 1.117099823 | 0.054550403 | 4.36E-06 | 1.58E-06 | 0.331747633 | 0.000334003 |
| NKTcells_CD8SP.PD1plus.CXCR3neg.PDL1plus | CD4TcellsTeff_Naive_TIGITplus | -0.374929721 | 0.622153254 | 0.098660571 | 0.997082975 | 0.049313315 | 0.000408064 | 7.21E-05 | 0.315974092 | 0.012659243 |
| NKTcells_CD16neg_CD95plus.CCR6neg.CXCR5plus | CD4TcellsTregs_Memory_CCR6plus.CD103plus.CD27plus | 0.3495635 | -0.706602027 | 0.045210604 | -1.056165527 | 0.068243543 | 2.64E-05 | 1.07E-05 | 0.368398755 | 0.001477906 |
| NKTcells_CD16neg.CD95plus.CCR6neg.CXCR5plus | CD4TcellsTregs_Memory_CD103plus.CD27plus | 0.345359326 | -0.804716092 | 0.006428154 | -1.150075418 | 0.071861143 | 2.45E-07 | 1.95E-07 | 0.377289727 | 3.10E-05 |
| NKTcells_CD16neg.CXCR5plus | CD4TcellsTregs_Memory_CD103plus.CD27plus | 0.172246726 | -0.762825061 | 0.079416529 | -0.935071787 | 0.380776554 | 2.37E-06 | 3.17E-05 | 0.742817584 | 0.000204443 |
| NKTcells_CD8SP_CD27plus.CD38plus.CD73plus | CD4TcellsTregs_Memory_CD103plus.CD27plus | 0.558730639 | -0.492150587 | 0.08506942 | -1.050881225 | 0.001998986 | 0.00780783 | 3.53E-05 | 0.049815724 | 0.104759939 |
| NKTcells_CD8SP_CD38plus.TIGITneg.CD73plus | CD4TcellsTregs_Memory_CD103plus.CD27plus | 0.572332882 | -0.482610776 | 0.079416529 | -1.054943658 | 0.001460215 | 0.009292741 | 3.15E-05 | 0.040618595 | 0.11698867 |
| NKTcells_CD16neg_CD95plus.CCR6neg.CXCR5plus | CD4TcellsTregs_Memory_CD103plus.CD62Lneg.CD27plus | 0.276825818 | -0.760617196 | 0.030490066 | -1.037443014 | 0.153843734 | 2.63E-06 | 5.84E-06 | 0.5243321 | 0.000223546 |
| NKTcells_CD16neg_CD95plus.CCR6neg.CXCR5plus | CD4TcellsTregs_Memory_CD103plus.CD62Lplus.CD27plus | 0.236642936 | -0.711385255 | 0.097365149 | -0.94802819 | 0.225352064 | 2.20E-05 | 6.35E-05 | 0.612162172 | 0.001269061 |
| NKTcells_CD16neg_CD161plus.KLRG1neg.CCR4plus | CD4TcellsTregs_Memory_CD103plus.HLADRneg.CD27plus | 0.340422636 | -0.678463654 | 0.077878654 | -1.018886289 | 0.076295689 | 7.24E-05 | 2.98E-05 | 0.387160271 | 0.003289146 |
| NKTcells_CD16neg_CD38plus.PD1plus.CCR4plus | CD4TcellsTregs_Memory_CD103plus.HLADRneg.CD27plus | 0.402766348 | -0.63785965 | 0.077878654 | -1.040625998 | 0.033585411 | 0.000260724 | 2.95E-05 | 0.260410937 | 0.008944813 |
| NKTcells_CD16neg_CD62Lplus.CD95plus.CXCR5plus | CD4TcellsTregs_Memory_CD103plus.HLADRneg.CD27plus | 0.397577907 | -0.697853922 | 0.030490066 | -1.095431829 | 0.036160724 | 3.66E-05 | 5.65E-06 | 0.270314382 | 0.001916672 |
| NKTcells_CD16neg_CD95neg.CXCR3neg.CD38plus | CD4TcellsTregs_Memory_CD103plus.HLADRneg.CD27plus | 0.563573536 | -0.577104124 | 0.026985013 | -1.140677661 | 0.001790247 | 0.001303691 | 4.59E-06 | 0.046389656 | 0.030485014 |
| NKTcells_CD16neg_CD95plus.CCR6neg.CXCR5plus | CD4TcellsTregs_Memory_CD103plus.HLADRneg.CD27plus | 0.477733849 | -0.823513611 | 0.000170576 | -1.30124746 | 0.010139058 | 7.35E-08 | 2.42E-09 | 0.135716278 | 1.12E-05 |
| NKTcells_CD16neg_CD95plus.ICOSplus | CD4TcellsTregs_Memory_CD103plus.HLADRneg.CD27plus | 0.511027634 | -0.51260287 | 0.097631177 | -1.023630504 | 0.005451858 | 0.005286051 | 6.43E-05 | 0.093778612 | 0.081131399 |
| NKTcells_CD16neg_CXCR5plus | CD4TcellsTregs_Memory_CD103plus.HLADRneg.CD27plus | 0.296373614 | -0.800590849 | 0.011790281 | -1.096964463 | 0.125669936 | 3.13E-07 | 6.69E-07 | 0.481040861 | 3.82E-05 |
| NKTcells_CD16neg_HLADRplus.CCR4plus | CD4TcellsTregs_Memory_CD103plus.HLADRneg.CD27plus | 0.173181769 | -0.802212661 | 0.030490066 | -0.975394429 | 0.378156827 | 2.84E-07 | 6.05E-06 | 0.740911357 | 3.52E-05 |
| NKTcells_CD16neg_PD1plus.CD161neg.CCR4plus | CD4TcellsTregs_Memory_CD103plus.HLADRneg.CD27plus | 0.366968168 | -0.653442108 | 0.085193324 | -1.020410275 | 0.054743515 | 0.000163051 | 3.74E-05 | 0.332396592 | 0.00620009 |
| NKTcells_CD16neg_TIGITplus.CXCR3neg.CCR4plus | CD4TcellsTregs_Memory_CD103plus.HLADRneg.CD27plus | 0.449460516 | -0.702606431 | 0.014798737 | -1.152066948 | 0.016416682 | 3.07E-05 | 1.62E-06 | 0.178016868 | 0.001668624 |
| NKTcells_CD8SP_CD27plus.CD38plus.CD73plus | CD4TcellsTregs_Memory_CD103plus.HLADRneg.CD27plus | 0.597014789 | -0.57589307 | 0.016831423 | -1.172907859 | 0.000796845 | 0.001341974 | 1.99E-06 | 0.027183573 | 0.031152967 |
| NKTcells_CD8SP_CD27plus.CD62Lplus.CD73plus | CD4TcellsTregs_Memory_CD103plus.HLADRneg.CD27plus | 0.45065941 | -0.585682473 | 0.085728628 | -1.036341884 | 0.016097196 | 0.001058614 | 4.33E-05 | 0.175963022 | 0.026183806 |
| NKTcells_CD8SP_CD27plus.CD73plus | CD4TcellsTregs_Memory_CD103plus.HLADRneg.CD27plus | 0.392696551 | -0.622773439 | 0.087970042 | -1.015469991 | 0.038725889 | 0.000401091 | 5.20E-05 | 0.279593645 | 0.012492396 |
| NKTcells_CD8SP_CD38plus.CCR4neg.CD73plus | CD4TcellsTregs_Memory_CD103plus.HLADRneg.CD27plus | 0.509159846 | -0.553255408 | 0.076153091 | -1.062415254 | 0.005654158 | 0.002259956 | 2.81E-05 | 0.095994107 | 0.045508219 |
| NKTcells_CD8SP_CD38plus.CD62Lplus.CD73plus | CD4TcellsTregs_Memory_CD103plus.HLADRneg.CD27plus | 0.534376521 | -0.512731041 | 0.085193324 | -1.047107562 | 0.003397224 | 0.005272751 | 3.94E-05 | 0.069817166 | 0.080997768 |
| NKTcells_CD8SP_CD38plus.CD73plus | CD4TcellsTregs_Memory_CD103plus.HLADRneg.CD27plus | 0.48709094 | -0.563054272 | 0.08506942 | -1.050145212 | 0.008568303 | 0.001811687 | 3.55E-05 | 0.122930393 | 0.038930302 |
| NKTcells_CD8SP_CD38plus.CD95neg.CD73plus | CD4TcellsTregs_Memory_CD103plus.HLADRneg.CD27plus | 0.447943739 | -0.577542181 | 0.089514915 | -1.025485921 | 0.016828366 | 0.001290079 | 5.49E-05 | 0.180693236 | 0.030253644 |
| NKTcells_CD8SP_CD38plus.TIGITneg.CD73plus | CD4TcellsTregs_Memory_CD103plus.HLADRneg.CD27plus | 0.591247636 | -0.562224379 | 0.022542099 | -1.153472016 | 0.000921979 | 0.001846414 | 3.29E-06 | 0.029997644 | 0.039484029 |
| NKTcells_CD8SP_CD73plus.CD62Lplus | CD4TcellsTregs_Memory_CD103plus.HLADRneg.CD27plus | 0.56818299 | -0.522939758 | 0.055318215 | -1.091122748 | 0.001609337 | 0.004300676 | 1.48E-05 | 0.043177213 | 0.070810554 |
| NKTcells_CD16neg_CD95plus.CCR6neg.CXCR5plus | CD4TcellsTregs_Memory_CD103plus.ICOSneg.CD27plus | 0.302761029 | -0.773367659 | 0.017229598 | -1.076128688 | 0.117341144 | 1.40E-06 | 2.12E-06 | 0.467259892 | 0.000131534 |
| NKTcells_CD16neg_CXCR5plus | CD4TcellsTregs_Memory_CD103plus.ICOSneg.CD27plus | 0.118859342 | -0.780412121 | 0.085193324 | -0.899271463 | 0.546900359 | 9.69E-07 | 3.76E-05 | 0.838762141 | 9.71E-05 |
| NKTcells_CD8SP_CD27plus.CD38plus.CD73plus | CD4TcellsTregs_Memory_CD103plus.ICOSneg.CD27plus | 0.530881174 | -0.492952479 | 0.097365149 | -1.023833653 | 0.003654256 | 0.00769269 | 6.34E-05 | 0.073041959 | 0.103820044 |
| NKTcells_CD8SP_CD38plus.TIGITneg.CD73plus | CD4TcellsTregs_Memory_CD103plus.ICOSneg.CD27plus | 0.544939816 | -0.47212357 | 0.098660571 | -1.017063386 | 0.002712276 | 0.011191513 | 7.07E-05 | 0.060341728 | 0.131905711 |
| NKTcells_CD16neg_CD95plus.CCR6neg.CXCR5plus | CD4TcellsTregs_Memory_CD103plus.TIGITneg.CD27plus | 0.198284972 | -0.726634116 | 0.098660571 | -0.924919088 | 0.311787951 | 1.19E-05 | 7.23E-05 | 0.691350231 | 0.000760469 |
| NKTcells_CD16neg_CD161plus.KLRG1neg.CCR4plus | CD4TcellsTregs_Memory_CD27plus | 0.337571342 | -0.67109849 | 0.085193324 | -1.008669832 | 0.078950891 | 9.27E-05 | 3.86E-05 | 0.392638856 | 0.003999215 |
| NKTcells_CD16neg_CD95plus.CCR6neg.CXCR5plus | CD4TcellsTregs_Memory_CD27plus | 0.32138302 | -0.658344418 | 0.098660571 | -0.979727437 | 0.095387089 | 0.000139896 | 7.17E-05 | 0.426959536 | 0.005510801 |
| CD8Tcells_Naive_ICOSplus.Ki67plus | CD8Tcells_Naive_Ki67plus | 0.758639391 | -0.135954991 | 0.097631177 | -0.894594382 | 2.89E-06 | 0.490307131 | 6.48E-05 | 0.000398445 | 0.808436452 |
| CD8Tcells_Naive_Eomesneg.Granzyme_Bplus | CD8Tcells_Naive_CD95plus.Heliosplus | 0.773956027 | -0.162549098 | 0.070223454 | -0.936505124 | 1.36E-06 | 0.40855954 | 2.42E-05 | 0.000216884 | 0.75810795 |
| CD8Tcells_Naive_Eomesneg.Tbetplus | CD8Tcells_Naive_CD95plus.Heliosplus | 0.81212699 | -0.225024899 | 0.014798737 | -1.037151889 | 1.55E-07 | 0.249620082 | 1.46E-06 | 3.58E-05 | 0.634471035 |
| CD8Tcells_Naive_Eomesneg.TNFaplus | CD8Tcells_Naive_CD95plus.Heliosplus | 0.800685509 | -0.351225326 | 0.006428154 | -1.151910835 | 3.11E-07 | 0.066853062 | 2.13E-07 | 6.36E-05 | 0.35492763 |
| CD8Tcells_Naive_IFNgneg.TNFaplus | CD8Tcells_Naive_CD95plus.Heliosplus | 0.761365785 | -0.195735113 | 0.069176067 | -0.957100898 | 2.54E-06 | 0.318177389 | 2.29E-05 | 0.000359764 | 0.69500935 |
| CD8Tcells_Naive_ICOSplus.Ki67plus | CD8Tcells_Naive_CD95plus.Tbetplus | 0.577819242 | -0.455014385 | 0.087970042 | -1.032833627 | 0.001281533 | 0.014979587 | 4.77E-05 | 0.03736259 | 0.15758981 |
| CD8Tcells_Naive_Ki67plus.Heliosplus | CD8Tcells_Naive_Eomesneg.TNFaplus | 0.566064588 | -0.503081312 | 0.070218099 | -1.0691459 | 0.001690413 | 0.006357167 | 2.38E-05 | 0.044628042 | 0.091466686 |

| | | | | | | | | | | |
|---|---|---|---|---|---|---|---|---|---|---|
| CD8Tcells_Naive_Eomesneg.TNFaplus | CD8Tcells_Naive_Eomesplus.Heliosplus | 0.59493856 | -0.417201052 | 0.097631177 | -1.012139612 | 0.000840072 | 0.027188482 | 6.51E-05 | 0.028181112 | 0.222638984 |
| CD8Tcells_Naive_ICOSplus.Ki67plus | CD8Tcells_Naive_Granzyme_Bplus | 0.649297291 | -0.378626556 | 0.083400141 | -1.027923847 | 0.000185206 | 0.046940959 | 3.39E-05 | 0.01004806 | 0.297351099 |
| CD8Tcells_Naive_ICOSplus.Ki67plus | CD8Tcells_Naive_Granzyme_Bplus.Eomesneg.Tbetplus | 0.346776621 | -0.719718624 | 0.032283081 | -1.066495246 | 0.070625454 | 1.58E-05 | 7.26E-06 | 0.374084985 | 0.000967921 |
| CD8Tcells_Naive_ICOSplus.Ki67plus | CD8Tcells_Naive_Granzyme_Bplus.Heliosneg.Tbetplus | 0.248969228 | -0.809837302 | 0.014257502 | -1.058806529 | 0.201394831 | 1.79E-07 | 1.05E-06 | 0.585630822 | 2.38E-05 |
| CD8Tcells_Naive_ICOSplus.Ki67plus | CD8Tcells_Naive_Granzyme_Bplus.IFNgneg.Tbetplus | 0.30646453 | -0.754872264 | 0.026403306 | -1.061336795 | 0.112702698 | 3.46E-06 | 4.24E-06 | 0.459493893 | 0.000277857 |
| CD8Tcells_Naive_ICOSplus.Ki67plus | CD8Tcells_Naive_Granzyme_Bplus.Tbetplus | 0.32153238 | -0.75670955 | 0.02161787 | -1.07824193 | 0.095224515 | 3.17E-06 | 2.96E-06 | 0.426657676 | 0.000257889 |
| CD8Tcells_Naive_ICOSplus.Ki67plus | CD8Tcells_Naive_ICOSplus.Granzyme_Bplus | 0.5173289 | -0.516679083 | 0.087970042 | -1.034007983 | 0.004814052 | 0.004876743 | 5.20E-05 | 0.086994622 | 0.076799178 |
| CD8Tcells_Naive_ICOSplus.Tbetplus | CD8Tcells_Naive_ICOSplus.Ki67plus | 0.549638244 | -0.479269026 | 0.09002755 | -1.028907269 | 0.002448018 | 0.009865962 | 5.57E-05 | 0.056481689 | 0.121571979 |
| CD8Tcells_Naive_TNFaplus.Tbetplus | CD8Tcells_Naive_ICOSplus.Ki67plus | 0.511997967 | -0.521929849 | 0.087970042 | -1.033927816 | 0.005349209 | 0.004389473 | 5.21E-05 | 0.092607714 | 0.071779718 |
| CD8Tcells_Naive_ICOSplus.Ki67plus | CD8Tcells_Naive_IFNgplus | 0.648493395 | -0.365475178 | 0.085728628 | -1.013968572 | 0.000189798 | 0.055812113 | 4.38E-05 | 0.010222368 | 0.324657643 |
| CD8Tcells_Naive_Tbetplus.TNFaplus | CD8Tcells_Naive_IFNgplus.Heliosplus | 0.502685035 | -0.530326045 | 0.087970042 | -1.033011079 | 0.006405457 | 0.003696569 | 5.28E-05 | 0.103547871 | 0.063912852 |
| CD8Tcells_Naive_ICOSplus.Ki67plus | CD8Tcells_Naive_IL21plus | 0.797440123 | -0.20917997 | 0.025838377 | -1.006620093 | 3.77E-07 | 0.28539158 | 4.03E-06 | 7.44E-05 | 0.667563304 |
| CD8Tcells_Naive_Tbetplus.TNFaplus | CD8Tcells_Naive_Ki67plus.Heliosplus | 0.226029578 | -0.7308528 | 0.085193324 | -0.956882378 | 0.247456224 | 1.00E-05 | 4.07E-05 | 0.634151533 | 0.00065806 |
| CD8Tcells_Naive_CD95plus.Heliosplus | CD8Tcells_Naive_TNFaplus | 0.66720358 | -0.308818853 | 0.098660571 | -0.976022433 | 0.000105327 | 0.109825582 | 6.97E-05 | 0.006648168 | 0.448642179 |
| CD8Tcells_Naive_Granzyme_Bneg.PD1plus | CD8Tcells_TPEX_CD27plus.CD95plus.CD62Lplus.PD1plus.TIGITplus | -0.074742357 | 0.795696616 | 0.085193324 | 0.870438973 | 0.705432713 | 4.16E-07 | 4.01E-05 | 0.909227078 | 4.84E-05 |
| CD8Tcells_Naive_Granzyme_Bneg.PD1plus | CD8Tcells_Naive_CD161plus.CD62Lneg.CCR6plus | 0.690369986 | -0.27943947 | 0.093687422 | -0.969809456 | 4.79E-05 | 0.14983597 | 5.93E-05 | 0.003672192 | 0.513603176 |
| CD8Tcells_Memory_CD95plus | CD8Tcells_Naive_KLRG1plus.CD161plus | 0.121053133 | -0.776999444 | 0.085193324 | -0.898052577 | 0.539471715 | 1.16E-06 | 4.15E-05 | 0.834786154 | 0.000112796 |
| CD8Tcells_Naive_Eomesneg.Granzyme_Bplus | BcellsnonPB_NonSwitchedMemory_CD27plus.IgDplus | 0.426529206 | -0.583642018 | 0.098660571 | -1.010171224 | 0.023608579 | 0.001112947 | 7.11E-05 | 0.217064262 | 0.027137469 |
| CD8Tcells_Naive_IL2plus.TNFaplus | BcellsnonPB_NonSwitchedMemory_CD27plus.IgDplus | 0.286167912 | -0.794876908 | 0.014257502 | -1.08104482 | 0.13986437 | 4.36E-07 | 1.09E-06 | 0.503501737 | 5.01E-05 |
| NKTcells_CD8SP_CD27plus.CD73plus | CD8Tcells_Naive_CD127plus | -0.051852033 | 0.798946005 | 0.087970042 | 0.850798038 | 0.79328912 | 3.45E-07 | 4.96E-05 | 0.939601888 | 4.16E-05 |
| NKTcells_CD8SP_CD62Lneg.CD73plus | CD8Tcells_Naive_CD127plus | -0.03006658 | 0.811390623 | 0.085193324 | 0.841457203 | 0.879283308 | 1.62E-07 | 4.04E-05 | 0.966126216 | 2.18E-05 |
| NKTcells_CD8SP_CD27plus.CD73plus | CD8Tcells_Naive_Eomesneg.CD127plus | -0.114240407 | 0.770424277 | 0.093687422 | 0.884664684 | 0.562694977 | 1.62E-06 | 5.90E-05 | 0.846521607 | 0.000148719 |
| NKTcells_CD8SP_CD62Lneg.CD73plus | CD8Tcells_Naive_Eomesneg.CD127plus | -0.088196563 | 0.776151436 | 0.098660571 | 0.864347999 | 0.655390473 | 1.21E-06 | 7.06E-05 | 0.889934405 | 0.000116801 |
| NKTcells_CD8SP_CD62Lplus.IFNgRplus.KLRG1plus | CD8Tcells_Naive_Granzyme_Bplus.Eomesneg.Tbetplus | -0.095171689 | 0.871885197 | 0.007775514 | 0.967056886 | 0.629992831 | 1.53E-09 | 3.81E-07 | 0.878770456 | 4.20E-07 |
| NKTcells_CD8SP_IFNgRplus.CCR6neg.KLRG1plus | CD8Tcells_Naive_Granzyme_Bplus.Eomesneg.Tbetplus | -0.147931119 | 0.76601836 | 0.085193324 | 0.913949479 | 0.452509883 | 2.02E-06 | 4.13E-05 | 0.787618699 | 0.00017961 |
| NKTcells_CD8SP_CD62Lplus.IFNgRplus.KLRG1plus | CD8Tcells_Naive_Granzyme_Bplus.Heliosneg.Tbetplus | -0.037057381 | 0.816925373 | 0.076153091 | 0.853982754 | 0.851493571 | 1.14E-07 | 2.81E-05 | 0.957791063 | 1.61E-05 |
| NKTcells_CD8SP_CD62Lplus.IFNgRplus.KLRG1plus | CD8Tcells_Naive_Granzyme_Bplus.IFNgneg.Tbetplus | -0.158398402 | 0.807994014 | 0.030490066 | 0.966392416 | 0.420788927 | 2.00E-07 | 5.93E-06 | 0.768951341 | 2.62E-05 |
| NKTcells_CD8SP_CD62Lplus.IFNgRplus.KLRG1plus | CD8Tcells_Naive_Granzyme_Bplus.Tbetplus | -0.132851441 | 0.802721944 | 0.046606434 | 0.935573385 | 0.50035572 | 2.76E-07 | 1.17E-05 | 0.813904009 | 3.43E-05 |
| NKTcells_CD16neg_CD161neg.KLRG1plus | CD8Tcells_Naive_ICOSplus.Ki67plus | 0.582857822 | -0.463882504 | 0.08506942 | -1.046740326 | 0.001134464 | 0.012901779 | 3.58E-05 | 0.034520968 | 0.143956013 |
| NKTcells_CD8SP_CD161neg.KLRG1plus | CD8Tcells_Naive_ICOSplus.Ki67plus | 0.51326648 | -0.512039696 | 0.096682972 | -1.025306176 | 0.005217494 | 0.005344832 | 6.21E-05 | 0.091269709 | 0.081784388 |
| NKTcells_CD8SP_CD38plus.CD62Lplus.CD73plus | CD8Tcells_Naive_ICOSplus.Ki67plus | -0.368873187 | 0.628939392 | 0.097631326 | 0.99781258 | 0.053403389 | 0.000337257 | 6.78E-05 | 0.328459281 | 0.01092058 |
| NKTcells_CD8SP_CD62Lplus.CCR6neg.KLRG1plus | CD8Tcells_Naive_ICOSplus.Ki67plus | 0.446564275 | -0.583923275 | 0.087970042 | -1.03048755 | 0.017210145 | 0.001105317 | 4.87E-05 | 0.182783821 | 0.027026167 |
| NKTcells_CD8SP_CD62Lplus.CD95neg.CD73plus | CD8Tcells_Naive_ICOSplus.Ki67plus | -0.411088123 | 0.648399858 | 0.062282975 | 1.059487981 | 0.029764697 | 0.000190339 | 1.90E-05 | 0.244896486 | 0.007025633 |
| NKTcells_CD8SP_CD62Lplus.CXCR3neg.CD73plus | CD8Tcells_Naive_ICOSplus.Ki67plus | -0.370081219 | 0.738511923 | 0.018258245 | 1.108593142 | 0.052567018 | 7.23E-06 | 2.33E-06 | 0.325762773 | 0.000504481 |
| NKTcells_CD4SP_HLADRplus.PD1plus.CD95plus | CD8Tcells_Naive_IL21plus.Granzyme_Bplus | -0.160920155 | 0.865633292 | 0.006428154 | 1.026553447 | 0.41333504 | 2.73E-09 | 1.75E-07 | 0.764141074 | 6.86E-07 |
| NKTcells_CD8SP_CD62Lplus.CXCR3neg.CD73plus | CD8Tcells_Naive_PD1plus.TNFaplus | 0.00725111 | -0.832555365 | 0.063936462 | -0.839806475 | 0.970787748 | 3.92E-08 | 2.09E-05 | 0.992560863 | 6.48E-06 |
| BcellsnonPB_NaiveTransitional_CD27neg.IgDplus.ICOSplus.CD | BcellsnonPB_DblNeg_CD27neg.IgDneg.CD38highplus | 0.625128204 | -0.445133444 | 0.062282975 | -1.070261648 | 0.00037556 | 0.017613659 | 1.83E-05 | 0.016464817 | 0.173418603 |
| Bcells_Plasmablasts_CXCR5plus.CD95plus | BcellsnonPB_DblNeg_CD27neg.IgDneg.IFNgRplus.CD73plus | 0.461045906 | -0.580392982 | 0.085193324 | -1.041438888 | 0.01353855 | 0.001204466 | 4.00E-05 | 0.159626889 | 0.028760983 |
| BcellsnonPB_NaiveTransitional_CD27neg.IgDplus.CD73plus.CD | BcellsnonPB_NaiveTransitional_CD27neg.IgDplus.CD11cplus | 0.349119546 | 0.972391416 | 8.26E-05 | 0.623271869 | 0.068618766 | 5.89E-18 | 3.90E-10 | 0.369424549 | 1.98E-14 |
| BcellsnonPB_SwitchedMemory_CD27plus.IgDneg.CD95plus | BcellsnonPB_NaiveTransitional_PBplus.CD27neg.IgDplus.CD95plus.CD25plus | 0.561847547 | -0.473183574 | 0.087970042 | -1.035031121 | 0.001862371 | 0.010985961 | 4.81E-05 | 0.047617053 | 0.130364503 |
| BcellsnonPB_SwitchedMemory_CD27plus.IgDneg.CD95plus | BcellsnonPB_NaiveTransitional_PBplus.CD27neg.IgDplus.IFNgRplus.CD25plus | 0.591444252 | -0.433881541 | 0.087970042 | -1.025325793 | 0.000917447 | 0.021066475 | 5.20E-05 | 0.029904305 | 0.192774507 |
| NKTcells_CD4SP_HLADRplus.PD1plus.CD95plus | NKTcells_CD8SP_CCR6neg.CD62Lneg.CD73plus | 0.251250331 | -0.747495131 | 0.055318215 | -0.998745463 | 0.197160857 | 4.85E-06 | 1.51E-05 | 0.5808417 | 0.000364678 |
| NKTcells_CD8SP_CD27plus.HLADRplus.KLRG1plus | NKTcells_CD8SP_CCR6neg.CD73plus | -0.272326952 | -0.892167108 | 0.087366396 | -0.619840156 | 0.1609208 | 1.83E-10 | 4.57E-05 | 0.534159905 | 6.87E-08 |
| NKTcells_CD8SP_CD62Lplus.CD95plus.KLRG1plus | NKTcells_CD8SP_CCR6neg.CD73plus | -0.039896805 | -0.828251572 | 0.088431369 | -0.788354767 | 0.840253799 | 5.31E-08 | 5.35E-05 | 0.954665142 | 8.46E-06 |
| NKTcells_CD4SP_HLADRplus.PD1plus.CD95plus | NKTcells_CD8SP_CCR6neg.CXCR3neg.CD73plus | 0.125742812 | -0.793195884 | 0.063046857 | -0.918938696 | 0.523752824 | 4.80E-07 | 2.00E-05 | 0.826824288 | 5.45E-05 |
| NKTcells_CD8SP_CD62Lplus.CD95plus.KLRG1plus | NKTcells_CD8SP_CCR6neg.KLRG1plus | 0.41732022 | 0.974151745 | 0.00011717 | 0.556831525 | 0.027140121 | 2.53E-18 | 1.11E-09 | 0.233596344 | 9.21E-15 |
| NKTcells_CD8SP_CD27plus.HLADRplus.KLRG1plus | NKTcells_CD8SP_CCR6neg.TIGITneg.CD73plus | -0.359169114 | -0.905719453 | 0.097631326 | -0.546550339 | 0.060507055 | 3.45E-11 | 6.74E-05 | 0.348458473 | 1.60E-08 |
| NKTcells_CD8SP_CD62Lplus.CD95plus.KLRG1plus | NKTcells_CD8SP_CCR6neg.TIGITneg.CD73plus | -0.054409554 | -0.833185277 | 0.087970042 | -0.778775723 | 0.783327363 | 3.75E-08 | 5.24E-05 | 0.936262389 | 6.26E-06 |
| NKTcells_CD4SP_HLADRplus.PD1plus.CD95plus | NKTcells_CD8SP_CD161neg.CD62Lneg.CD73plus | 0.263365813 | -0.797915547 | 0.014798737 | -1.061281359 | 0.175700565 | 3.66E-07 | 1.45E-06 | 0.554371888 | 4.37E-05 |
| NKTcells_CD4SP_HLADRplus.PD1plus.CD95plus | NKTcells_CD8SP_CD161neg.CXCR3neg.CD73plus | 0.119582288 | -0.819611275 | 0.031807925 | -0.939193563 | 0.544447055 | 9.55E-08 | 6.46E-06 | 0.837550822 | 1.39E-05 |
| NKTcells_CD8SP_TIGITplus.CD73plus | NKTcells_CD8SP_CD27plus.CD73plus | 0.091892931 | 0.900133823 | 0.014257502 | 0.808240891 | 0.641882364 | 7.06E-11 | 1.05E-06 | 0.884224426 | 3.00E-08 |
| NKTcells_CD8SP_CD38plus.CCR4neg.CD73plus | NKTcells_CD8SP_CD27plus.HLADRplus.KLRG1plus | -0.362346593 | -0.909560207 | 0.087970042 | -0.547213614 | 0.058104346 | 2.06E-11 | 5.13E-05 | 0.341681635 | 1.02E-08 |
| NKTcells_CD8SP_CD38plus.CD73plus | NKTcells_CD8SP_CD27plus.HLADRplus.KLRG1plus | -0.352328153 | -0.908037576 | 0.087970042 | -0.555709423 | 0.065942488 | 2.53E-11 | 4.92E-05 | 0.362534611 | 1.22E-08 |
| NKTcells_CD16neg_IFNgRplus.CCR6neg.KLRG1plus | NKTcells_CD8SP_CD38plus.CD62Lplus.CD73plus | -0.031536196 | -0.852966968 | 0.049176473 | -0.821430772 | 0.87342856 | 8.17E-09 | 1.26E-05 | 0.96447514 | 1.74E-06 |
| NKTcells_CD4SP_HLADRplus.PD1plus.CD95plus | NKTcells_CD8SP_CD38plus.CXCR3neg.CD73plus | 0.129660478 | -0.814035808 | 0.032283081 | -0.943696287 | 0.510793387 | 1.37E-07 | 7.20E-06 | 0.819708777 | 1.90E-05 |
| NKTcells_CD4SP_HLADRplus.PD1plus.CD95plus | NKTcells_CD8SP_CD62Lneg.CD73plus | 0.280318368 | -0.733208086 | 0.055318215 | -1.013526453 | 0.14850526 | 9.08E-06 | 1.52E-05 | 0.516462468 | 0.000606707 |
| NKTcells_CD4SP_HLADRplus.PD1plus.CD95plus | NKTcells_CD8SP_CD62Lneg.CD95neg.CD73plus | 0.331050966 | -0.724472448 | 0.035667919 | -1.055523413 | 0.085288159 | 1.31E-05 | 8.26E-06 | 0.406360452 | 0.000821151 |
| NKTcells_CD16neg_IFNgRplus.CCR6neg.KLRG1plus | NKTcells_CD8SP_CD62Lplus.CCR6neg.KLRG1plus | 0.309133621 | 0.899160418 | 0.087970042 | 0.590026797 | 0.109445097 | 7.97E-11 | 4.92E-05 | 0.453622239 | 3.34E-08 |
| NKTcells_CD16neg_IFNgRplus.CCR6neg.KLRG1plus | NKTcells_CD8SP_CD62Lplus.CD95neg.CD73plus | -0.145475758 | -0.86398839 | 0.085193324 | -0.718512631 | 0.460130307 | 3.17E-09 | 3.96E-05 | 0.792101779 | 7.80E-07 |
| NKTcells_CD8SP_CCR6neg.CD73plus | NKTcells_CD8SP_KLRG1plus | -0.068649414 | -0.841726142 | 0.085728628 | -0.773076728 | 0.728511164 | 1.99E-08 | 4.22E-05 | 0.917940331 | 3.67E-06 |
| NKTcells_CD8SP_CD38plus.CCR4neg.CD73plus | NKTcells_CD8SP_KLRG1plus | -0.111009484 | -0.848488241 | 0.0902048 | -0.737478757 | 0.573864825 | 1.18E-08 | 5.63E-05 | 0.852039449 | 2.34E-06 |
| NKTcells_CD8SP_CD38plus.CD73plus | NKTcells_CD8SP_KLRG1plus | -0.123101501 | -0.848874786 | 0.097631326 | -0.725773285 | 0.532578731 | 1.14E-08 | 6.62E-05 | 0.831546969 | 2.29E-06 |
| NKTcells_CD8SP_CD62Lplus.CCR6neg.KLRG1plus | NKTcells_CD8SP_KLRG1plus | 0.303853337 | 0.899857318 | 0.085728628 | 0.596003981 | 0.115958676 | 7.31E-11 | 4.26E-05 | 0.464870552 | 3.09E-08 |
| NKTcells_CD4SP_HLADRplus.PD1plus.CD95plus | NKTcells_CD8SP_PD1plus.CCR6neg.CD73plus | 0.230644175 | -0.736855743 | 0.079416529 | -0.967499917 | 0.237676412 | 7.76E-06 | 3.09E-05 | 0.624303264 | 0.000533853 |
| NKTcells_CD4SP_HLADRplus.PD1plus.CD95plus | NKTcells_CD8SP_PD1plus.CD161neg.CD73plus | 0.257537063 | -0.71734693 | 0.085193324 | -0.974883993 | 0.185810552 | 1.74E-05 | 3.77E-05 | 0.567418659 | 0.001051391 |

Table S2 Reagents and Resources

| REAGENT or RESOURCE | SOURCE | IDENTIFIER |
|---|---|---|
| Antibodies | | |
| CD45 89Y HI30 | Fluidigm | Cat# 3089003<br>AB_2661851 |
| CD19 Cd113 HIB19 | Biolegend | Cat#302201<br>AB_314231 |
| CD3 Cd116 UCHT1 | Biolegend | Cat#300401<br>AB_314055 |
| TCRgd 141Pr 11F2 | Check with AW | Cat# 3152008B<br>AB_2687643 |
| CD45RA 143Nd | Fluidigm | Cat# 3143006B<br>AB_2651156 |
| CD38 144Nd HIT2 | Fluidigm | Cat# 3144014B<br>AB_2687640 |
| CD4 145Nd RPA-T4 | Fluidigm | Cat# 3145001<br>AB_2661789 |
| CD8a 146Nd RPA-T8 | Fluidigm | Cat# 3146001<br>AB_2687641 |
| CD161 147Sm HP-3G10 | Biolegend | Cat#339902<br>AB_1501090 |
| CD127 149Sm A019D5 | Fluidigm | Cat# 3149011<br>AB_2661792 |
| CD14 151Eu M5E2 | Fluidigm | Cat# 3151009B<br>AB_2810244 |
| CD95 152Sm DX2 | Fluidigm | Cat# **3152017B** |
| HLA-DR 153Eu | Biolegend | Cat#307602<br>AB_314680 |
| CD11c 159Tb Bu15 | Fluidigm | Cat#3159001<br>AB_2661800 |
| IgD 162Dy IA6-2 | Biolegend | Cat#348202<br>AB_10550095 |
| ICOS 163Dy C398.4A | Biolegend | Cat#313541 AB_2728259 |
| CD123 165Ho | Biolegend | Cat#306002<br>AB_314576 |
| CD56 166Er NCAM16.2 | BD Biosciences | Cat# 559043<br>AB_397180 |
| CD27 167Er M-T271 | Biolegend | Cat#356401 AB_2561786 |
| CD25 169Tm 2A3 | Fluidigm | Cat# 3169003<br>AB_2661806 |
| CXCR5 170Er | BD Bioscience | Cat#552032 AB_394324 |
| CD19 174Yb HIB19 | Biolegend | Cat#302201<br>AB_314231 |

| | | |
|---|---|---|
| PD-1 175Lu EH12.2H7 | Fluidigm | Cat# 3175008<br>AB_2687629 |
| CD16 209Bi 3G8 | Fluidigm | Cat# 3209002B<br>AB_2756431 |
| IL-4 142 MP4-25D2 | Fluidigm | Cat# **3142002B** |
| Helios 148Nd 22F6 | Biolegend | Cat#137202<br>AB_10900638 |
| IL-22 150Nd 22URTI | Fluidigm | Cat# 3150007B<br>AB_2810972 |
| Granzyme K 154Sm | Biolegend | Cat#370502<br>AB_2566673 |
| Ki-67 155Gd Ki-67 | Biolegend | Cat#350501<br>AB_10662749 |
| Tbet 156Gd | Biolegend | Cat#644801<br>AB_1595608 |
| IL-2 158Gd Bu15 | Fluidigm | Cat# 3158007B<br>AB_2864735 |
| IL-6 160Gd | Biolegend | Cat#501101<br>AB_315149 |
| Il-17A 161Dy BL168 | Fluidigm | Cat# **3161008B** |
| Eomes 164Dy WD1928 | eBioscience | Cat#14-4877-82<br>AB_2572882 |
| IFNg 168Er B27 | Fluidigm | Cat# **3168005B** |
| Granzyme B 171Yb BG11 | Fluidigm | Cat# 3171002B<br>AB_2687652 |
| IL-21 172Yb 3A3-Ns | Fluidigm | Cat# 3172011B,<br>AB_2810975 |
| FoxP3 173Yb 259D/C7 | Biolegend | Cat# 320201<br>AB_430884 |
| TNFa 176Yb Mab11 | Biolegend | Cat#502901 AB_315253 |
| CD103 $142^{Nd}$ Ber-ACT8 | Biolegend | Cat#350202 AB_10639864 |
| PD-L1 148Nd 29E.2A3 | Fluidigm | Cat#3148017B |
| CXCR3 150Nd G025H7 | Biolegend | Cat#353702<br>AB_10983073 |
| CCR6 154Sm G034E3 | Biolegend | Cat#353427<br>AB_2563725 |
| CD62L 155Gd DREG-56 | Biolegend | Cat#304802<br>AB_314462 |
| Siglec-1 156Gd 7-239 | Biolegend | Cat#346002<br>AB_2189031 |
| CCR4 158Gd L291H4 | Fluidigm | Cat#3158032A |
| CD39 160 Gd A1 | Fluidigm | Cat# 3160004B, AB_2687648 |
| CD49b 161Dy P1E6-C5 | Fluidigm | Cat#3161012B |
| TIGIT 164Dy MBSA43 | Thermo Fisher | Cat# 16-9500-82<br>AB_10718831 |

| | | |
|---|---|---|
| CD45RO 165Ho UCHL1 | Fluidigm | 3165011B<br>AB_2756423 |
| CD73 168 Er AD2 | Fluidigm | Cat# 3168015B<br>AB_2810249 |
| IFNgR 171Yb | BD Bioscience | Cat# 558935<br>AB_397164 |
| LAG3 172 Yb 874501 | R&D systems | Cat# MAB23193-SP |
| CRTH2 173Yb BM16 | Biolegend | Cat#350102 AB_10639863 |
| KLRG1 174Yb | Biolegend | Cat#368602<br>AB_2566256 |
| CD123 176Yb | Biolegend | Cat#306002<br>AB_314576 |
| pSTAT1 (Y701)-AF488 4a | BD Bioscience | Cat#612596<br>AB_399879 |
| pSTAT3 (Y705)-PE 4/P-STAT3 | BD Bioscience | Cat#612569<br>AB_399860 |
| pSTAT5 (Y694)-PE-Cy7 47/Stat5 | BD Bioscience | Cat#560117<br>AB_1645546 |
| Mouse anti-Total Stat1 PE (N-Terminus) | BD Bioscience | Cat#558537<br>AB_647231 |
| Mouse anti-Stat3 APC M59-50 | BD Bioscience | Cat#560392<br>AB_1645463 |
| Stat1 Phospho (Ser727) PE/Cyanine7 A15158B | Biolegend | Cat#686407<br>AB_2650781 |
| STAT3 Phospho (Ser727) PerCP A16089B | Biolegend | Cat#698909<br>AB_2734541 |
| CD4 SB780 SK3 | Thermo Fisher | Cat#78-0047-42<br>AB_2744896 |
| CD4 BV650 SK3 | BD Bioscience | Cat#563875 |
| CD4 BV510 OKT4 | Biolegend | Cat#317443<br>AB_2561377 |
| CD8a BV711 SK1 | Biolegend | Cat#344733<br>AB_2565242 |
| CD8a BV650 SK1 | Biolegend | Cat#344741<br>AB_2566512 |
| CD3 BV605 OKT3 | Biolegend | Cat#317321<br>AB_11126166 |
| CD27 BUV395 M-T271 | BD Bioscience | Cat#740291<br>AB_2740030 |

| | | |
|---|---|---|
| CD45RA BUV737 HI100 | BD Bioscience | Cat#612846<br>AB_2870168 |
| CD45RA BV711 HI100 | BD Bioscience | 563733 |
| CD3 BV605 OKT3 | Biolegend | Cat#317321<br>AB_11126166 |
| CD62L BV510 DREG-56 | Biolegend | Cat#304843<br>AB_2617002 |
| CD56 BV785 5.1H11 | Biolegend | Cat#362549<br>AB_2566058 |
| CXCR5 BB700 RF8B2 | BD Bioscience | Cat#566470<br>AB_2738392 |
| CCR6 BV421 G034E3 | Biolegend | Cat#353407<br>AB_10916530 |
| LIVE/DEAD Fixable Aqua Dead Cell Stain Kit | Invitrogen | Cat#L34966 |
| Cisplatin | Enzo Life Sciences | Cat# ALX-400-040-M250 |
| CD38 PerCP/Cy5.5 HIT2 | Biolegend | Cat#303522<br>AB_893314 |
| CD39 BV711 TU66 | BD Bioscience | Cat#563680<br>AB_2738369 |
| CD8a AF700 RPA-T8 | Biolegend | Cat#301028<br>AB_493745 |
| CD45RA FITC HI100 | Biolegend | Cat#304106<br>AB_314410 |
| CD62L BV510 DREG-56 | Biolegend | Cat#304844<br>AB_2617003) |
| CD27 BUV395 L128 | BD Bioscience | Cat#563815<br>AB_2744349 |
| TIGIT PE eFluor610 MBSA43 | Invitrogen | Cat#61-9500-42<br>AB_2723715 |
| CXCR3 AF647 G025H7 | Biolegend | Cat#353712 AB_10962948 |
| PD-1 PE-Cy7 EH122H7 | Biolegend | Cat#329918<br>AB_2159324 |
| CD73 PE AD2 | Biolegend | Cat#344004<br>AB_2298698 |
| CD95 BV421 DX2 | Biolegend | Cat#305624AB_2561830 |
| Biological Samples | | |
| Human serum | BRI Repository | https://www.benaroyaresearch.org/what-is-bri/scientists-and-laboratories/core-labs/translational-core-laboratory |
| Human peripheral mononuclear cells | BRI Repository | https://www.benaroyaresearch.org/what-is-bri/scientists-and- |
| Chemicals, Peptides, and Recombinant Proteins | | |
| Human Interferon alpha | PBL | Cat#11101-1 |

| | | |
|---|---|---|
| Human IL-6 | BD Pharmingen | Cat#550071 |
| Human Interferon Beta | PBL | Cat#11415-1 |
| Human IL-2 | Peprotech | Cat# 200-02 |
| Human Interferon gamma | R&D systems | Cat# 285-IF-100 |
| Critical Commercial Assays | | |
| V-Plex Th17 Panel 1 | Mesoscale Discovery | Cat# K15085D-1 |
| V-Plex Proinflammatory Panel 1 | Mesoscale Discovery | Cat# K15049D-1 |
| Pan T cell Isolation kit, human | Miltenyi Biotec | Cat# 130-096-535 |
| Cytomegalovirus IgG Human ELISA Kit | Abcam | Cat#ab108639 |
| Maxpar Fix and Perm Buffer | Fluidigm | Cat#201067 |
| Maxpar X8 Antibody Labeling Kit 160 Gd | Fluidigm | Cat#201160A |
| Maxpar MCP9 Antibody Labeling Kit 116Cd | Fluidigm | Cat#201116A |
| MaxPar Nuclear Antigen Staining Buffer Set | Fluidigm | Cat# 201063 |
| Perm Buffer III | BD Phosflow | Cat#558050 |
| Cytofix | BD Bioscience | Cat#554722 |
| Fix buffer 1 | BD Biosciences | Cat#557870 |
| Permeabilization Buffer | eBioscience | Cat#00-8333-56 |
